# Potent SARS-CoV-2 binding and neutralization through maturation of iconic SARS-CoV-1 antibodies

**DOI:** 10.1101/2020.12.14.422791

**Authors:** Romain Rouet, Ohan Mazigi, Gregory J. Walker, David B. Langley, Meghna Sobti, Peter Schofield, Helen Lenthall, Jennifer Jackson, Stephanie Ubiparipovic, Jake Y. Henry, Arunasingam Abayasingam, Deborah Burnett, Anthony Kelleher, Robert Brink, Rowena A. Bull, Stuart Turville, Alastair G. Stewart, Christopher C. Goodnow, William D. Rawlinson, Daniel Christ

## Abstract

Antibodies against coronavirus spike protein potently protect against infection and disease, however it remains unclear if such protection can be extended to variant coronaviruses. This is exemplified by a set of iconic and well-characterized monoclonal antibodies developed after the 2003 SARS outbreak including mAbs m396, CR3022, CR3014 and 80R, which potently neutralize SARS-CoV-1, but not SARS-CoV-2. Here we explore antibody maturation strategies to change and broaden their specificity, enabling potent binding and neutralization of SARS-CoV-2. Using targeted mutagenesis as well as light chain shuffling on phage, we identified variants with considerably increased affinity and neutralization potential. The most potent antibody, derived from the NIH-developed mAb m396, neutralized live SARS-CoV-2 virus with a half-maximal inhibitory concentration (IC_50_) of 160 ng/ml. Intriguingly, while many of the matured clones maintained specificity of the parental antibody, new specificities were also observed, which was further confirmed by X-ray crystallography and cryo-electron microscopy, indicating that a limited set of antibodies can give rise to variants targeting diverse epitopes. Our findings open up over 15 years of antibody development efforts against SARS-CoV-1 to the SARS-CoV-2 field and outline general principles for the maturation of antibody specificity against emerging viruses.

## INTRODUCTION

The emergence of at least three coronaviruses (SARS-CoV-1, SARS-CoV-2 and MERS) in the human population in the last two decades has highlighted the need for rapid and sustained development of prophylactic and therapeutic modalities. Among such modalities, antibody reagents blocking the interaction of the viral spike protein with human receptor (angiotensin converting enzyme 2 (ACE2) in the case of SARS-CoV-1 and CoV-2, and dipeptidyl peptidase 4 (DPP4) in the case of MERS) are among the most promising^1–3^. Various approaches have been used to identify neutralizing antibodies, including the identification of B-cells from convalescent patients ^4,5^, the immunization of humanized transgenic mice ^6^, or through the use of *in vitro* library display approaches against viral spike protein (or more commonly its receptor binding domain (RBD))^7,8^.

Here we employed a different approach based on the re-engineering and maturation of previously reported antibodies against SARS-CoV-1. Although such antibodies generally do not bind and neutralize SARS-CoV-2, we speculated that the relatively high level of sequence identify of the RBD of the two viruses (76% amino acid identity ^9,10^) would allow us to shift antibody specificity through limited changes in antibody variable regions.

We focused our attention on four well-characterized monoclonal antibodies (m396 ^11^, CR3022 ^12^, CR3014 ^13^ and 80R ^14^) which bind and neutralize SARS-CoV-1 with equilibrium binding and IC_50_ constants in the nanomolar range. Crystal structures have been reported for m396 ^11^, CR3022 ^12^ and 80R ^14^ in complex with RBD; these reveal binding to a diverse set of epitopes, with m396 and 80R binding to distinct, but adjacent, epitopes overlapping with the ACE2 binding site (Fig. 1a). Although no structural information has been reported for CR3014, the antibody has been shown to block ACE2 binding ^12^. In marked contrast, CR3022 binds to an epitope distant to the ACE2 binding site which is largely conserved between SARS-CoV-1 and SARS-CoV-2 ^15^. Unlike m396, CR3014 and 80R, CR3022 displays residual binding to SARS-CoV-2 RBD; however, it does not detectably neutralize live SARS-CoV-2 virus ^15^.

**Figure 1.**
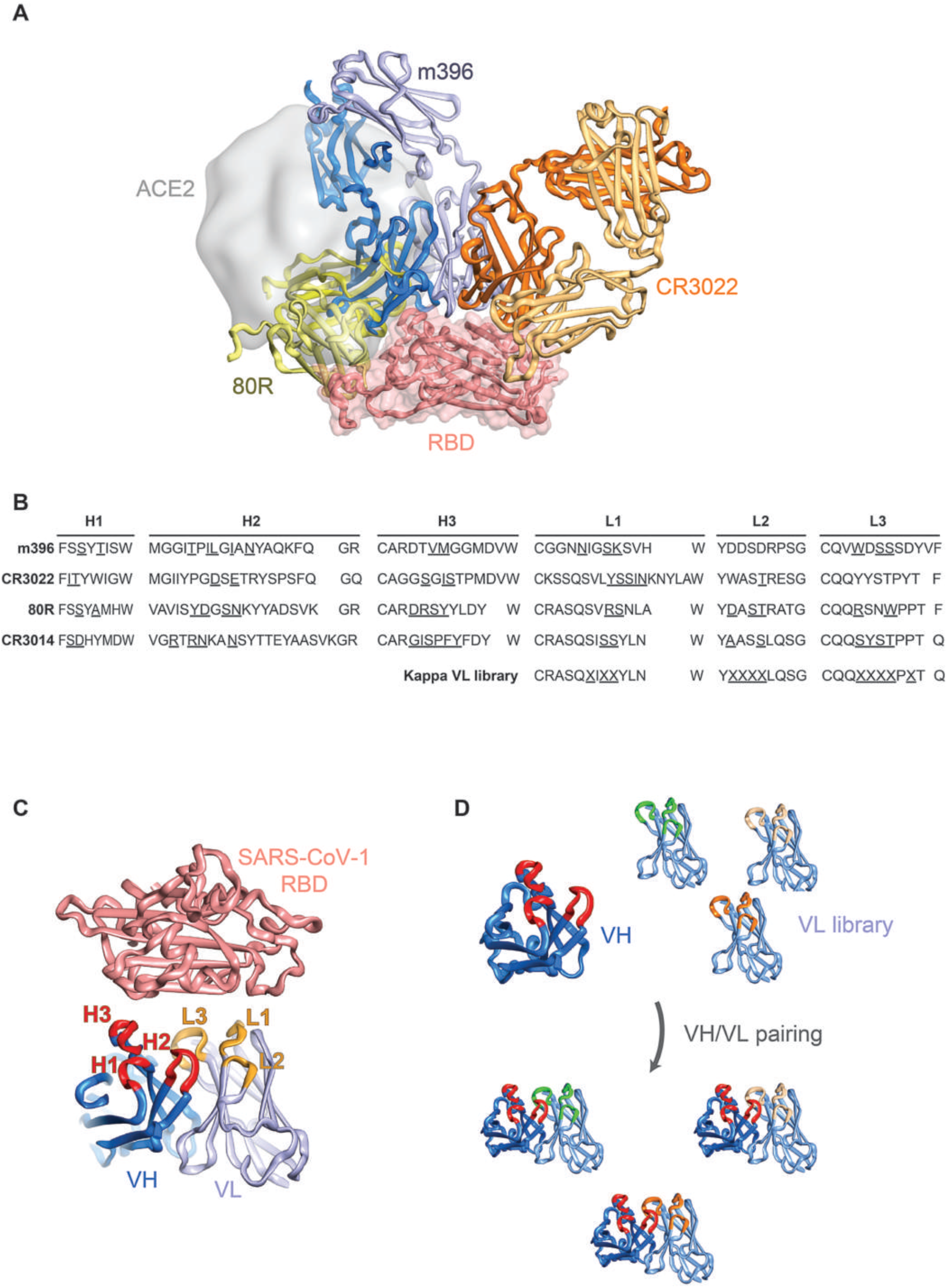
Design of antibody libraries. **(A)** Structures of SARS-CoV-1 antibodies superposed on the surface of RBD (salmon) with ACE2 highlighted in grey. (**B)** CDR regions of SARS-CoV-1 antibodies with randomized position underlined. (**C)** Site-directed mutagenesis strategy with targeted antibody CDR regions highlighted (VH in red, V_L_ in orange). (**D)** Light chain shuffling strategy with variant kappa V_L_ domains highlighted.

For the re-engineering strategy, we focused on two well-established *in vitro* methods for antibody affinity maturation: (I) site-directed mutagenesis of complementarity determining regions (CDR) of human variable domains ^16^ and (II) light chain shuffling ^17^ (Fig. 1B-D). Library design based on the reported structures of m396, CR3022 and 80R in complex with RBD was used for the construction of site-directed mutagenesis repertoires, with antibody contact residues with antigen targeted for diversification. For the alternative light chain shuffling approach, a previously described highly diverse synthetic antibody library based on a single Vκ1 framework was utilized ^18,19^. Both library classes were then selected for binding to SARS-CoV-2 RBD using iterative selections on biotinylated antigen (100 nM to 500 pM range). Using these approaches, we rapidly identified human antibody variants with potent affinity and neutralization potential for SARS-CoV-2.

## RESULTS

### Generation and selection of SARS-CoV-2 binding antibodies by site-directed mutagenesis

For the design of site-directed mutagenesis libraries, we utilized previously reported crystal structures of antibodies developed against SARS-CoV-1 in complex with either cognate RBD (80R – PDB ID 2ghw ^20^; m396 - PDB ID 2dd8 ^21^) or SARS-CoV-2 RBD in the case of the cross-specific CR3022 antibody (PDB ID 6w41 ^15^). Based on the structures, we selected contact and proximal residues in CDR regions of V_H_ and V_L_, which were targeted for diversification by Kunkel mutagenesis ^16^ (Fig. 1A-C and Supplementary Table 1; all six CDR regions were targeted for CR3014 for which no structural information has been reported). Library construction was carried out in an antibody single-chain Fv (scFv) format, resulting in 6.1 × 10^8^, 2.3 × 10^7^, 3.4 × 10^7^ and 5.7 × 10^7^ clones for m396, CR3022, CR3014 and 80R respectively. We performed four rounds of phage display selection, using decreasing amounts of SARS-CoV-2 RBD for selection (see Methods); this resulted in enrichment of SARS-CoV-2 specific binders for the libraries (with the exception of CR3014), as indicated by polyclonal phage ELISA (Fig. 2A). Screening of individual clones by monoclonal soluble ELISA was performed after round 4, followed by sequencing and cloning of non-redundant variants into an IgG expression vector. After production in CHO cells, monoclonal antibodies were characterized for binding to recombinant RBD by biolayer-interferometry (BLI) and for neutralization of live SARS-CoV-2 virus in Vero E6 cells.

**Figure 2.**
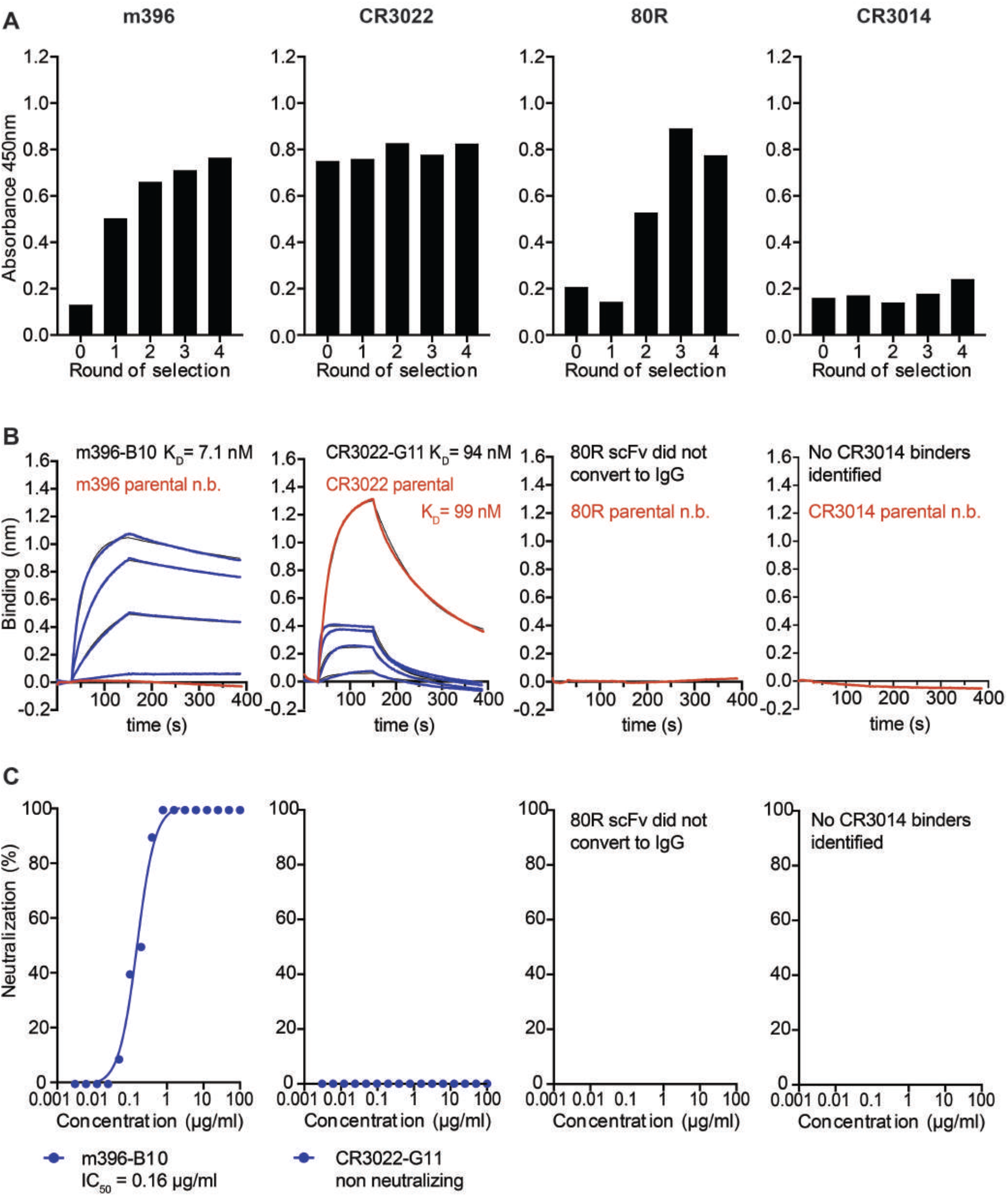
Selection of SARS-CoV-2 specific antibodies from site-directed mutagenesis libraries of SARS-CoV-1 binders. (**A**) Enrichment of scFv antibody binders by phage display (polyclonal phage ELISA). (**B)** Biolayer interferometry affinity measurements of soluble SARS-CoV-2 RBD (at 400 nM, 200 nM, 100 nM, 50 nM; highest concentration only shown for parental antibodies) binding to immobilized antibody (see Methods). (**C)** Neutralization of live SARS-CoV-2 virus in Vero E6 cells (IgG format).

In the case of m396, two variants (designated B10 and C4) were chosen for further characterization, with both antibody variants encoding several mutations in V_H_ and V_L_ (Supplementary Sequences). Both variants displayed high monovalent binding affinity to soluble SARS-CoV-2 RBD with equilibrium binding constants (K_D_) in the low nanomolar range (7.1 nM in the case of B10 and 13 nM in the case of C4) (Fig. 2B, Supplementary Fig. 1A and Table 1). Both variants also potently neutralized live SARS-CoV-2 virus with IC_50_s of 160 ng/ml and 340 ng/ml, respectively (Fig. 2C and Supplementary Fig. 1E). In addition to live virus, m396-B10 also potently neutralized both SARS-CoV-1 and SARS-CoV-2 pseudoparticles (with IC_50_s of 2.2 and 0.3 μg/ml, respectively) (Supplementary Fig. 1L).

**Table 1.**
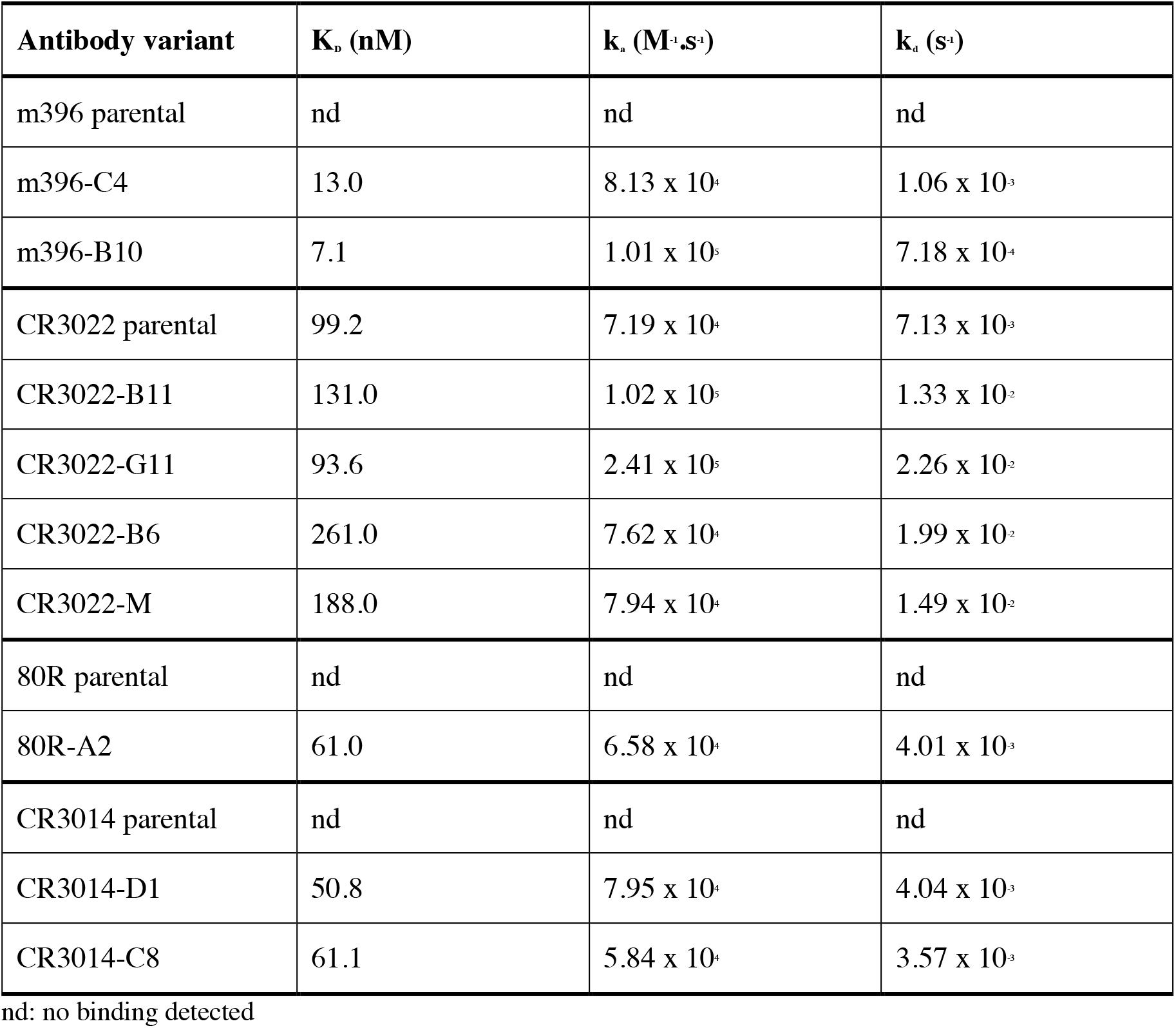
Affinity of monoclonal antibodies (biolayer interferometry)

The B10 variant of m396 was further selected for structural characterization by crystallography in a Fab format, both in isolation and in complex with SARS-CoV-2 RBD. Crystals were obtained for the unliganded B10 Fab, which diffracted to 2.3 Å (Supplementary Table 2). Although no crystals were obtained for the B10 complex, analysis of the structure of the m396 parent bound to SARS-CoV-1 RBD (PDB ID 2dd8 – Supplementary Fig. 2)^21^ reveals that the bulk of the contact surface is contributed by heavy chain interactions (517 Å^2^ buried surface vs 370 Å^2^ for the light chain) in which CDR H1 and H2 line one side of a cleft, whilst H3 lines the other side, into which a loop of CoV1 RBD projects (residues 484-492 SARS-CoV-1 numbering, residues 498-506 SARS-CoV-2 numbering). The m396-B10 clone contains several heavy chain CDR mutations relative to the parental m396 antibody: two in H1, four in H2, and two in H3 (Supplementary Information). Although the overall RBD fold is conserved between SARS-CoV-1 and SARS-CoV-2 (Supplementary Fig. 2, colored in yellow and salmon respectively), the loop bound by the m396 heavy chain cleft comprises a local divergence hotspot containing multiple substitutions: Y498Q, T499P, T501N, I503V (all SARS-CoV-2 numbering), considerably more divergent than the overall RBD. The crystal structure of the m396-B10 Fab described here lacks electron density at most of these CDR positions, indicating conformational plasticity in the unliganded state. In contrast to heavy chain, light chain residues form more limited contacts in the m396 parental complex, with CDR L1 and L3 contacting a surface with considerably greater conservation between SARS-CoV-1 and SARS-CoV-2. A total of two mutations were selected in the m396-B10 L3 region (Supplementary Sequences) which contacts the RBD in the parental m396 complex, with no mutations observed in L1 and L2 regions.

To further define the epitope of m396 B10, we carried out epitope binning by BLI, which indicated competition with recombinant ACE2 (Supplementary Fig. 3A). In addition, we generated a triple mutant within the ACE2 binding site of SARS-CoV-2 RBD (comprising T500A, N501A and Y505A, SARS-CoV-2 numbering), predicted to interfere with parental m396 binding. Mutation of this region in RBD resulted in complete loss of binding of m396-B10 (as well as for m396-C4) (Supplementary Fig. 3B), suggesting that these variants bind to an epitope within the ACE2 binding site, as previously demonstrated for the parental m396 SARS-CoV-1 RBD interaction ^11^.

In addition to m396, we selected two variants of CR3022 for further characterization (clones G11 and B11). When expressed in an IgG format both antibodies displayed similar equilibrium binding constants (K_D_) for SARS-CoV-2 RBD as the parental CR3022 antibody: 94 nM for B11 and 131 nM for G11, respectively, compared to 99 nM for CR3022 (G11: k_a_2.4×10^5^ M^−1^.s^−1^, k_d_2.2 × 10^−2^ s^−1^ ; B11: k_a_1.0×10^5^ M^−1^.s^−1^, k_d_1.3 × 10^−2^s^−1^ ; CR3022 parental: k_a_ 7.2×10^4^ M^−1^.s^−1^, k_d_ 7.1 × 10^−3^ s^−1^) (Fig. 2B, Table 1 and Supplementary Fig. 1A). However, both the parental CR3022 IgG, as well as the G11 and B11 variants, did not detectably neutralize SARS-CoV-2 virus (Fig. 2C and Supplementary Fig. 1E) ^15^.

In contrast to the m396 and CR3022 phage display selections, no enrichment was observed for the selection of the CR3014 site-directed mutagenesis library (Fig. 2A). In the case of the 80R selection, polyclonal ELISA indicated the selection of binders, which could also be detected by soluble ELISA in scFv format (Supplementary Fig. 4). Three of the selected clones, designated C9, D1 and D10 were converted into an IgG format, however binding was lost upon antibody format conversion and the clones did not display detectable neutralization activity (Fig. 2B-C and Supplementary Fig. 1E).

### Generation and selection of SARS-CoV-2 binding antibodies by light chain shuffling

In addition to the site-directed mutagenesis approach described as above, we investigated light chain shuffling as a strategy for shifting the specificity of antibodies from SARS-CoV-1 towards SARS-CoV-2 ^17^. We utilized splice overlap extension PCR ^22^ to pair DNA encoding variable heavy domains of each of the four antibodies analyzed here (m396, CR3022, CR3014 and 80R) with a kappa light chain library on phage in a scFv format (Fig. 1D). The synthetic light chain library is based on the human Vκ1 framework, with diversity introduced at CDR L1, L2 and L3 position ^18^. After ligation and electroporation into *E. coli* TG1, light chain shuffled libraries of 5 × 10^7^, 1 × 10^8^, 4 × 10^7^ and 1 × 10^8^ clones were obtained for m396, CR3022, CR3014 and 80R, respectively. Three rounds of phage display selection were performed using decreasing amounts of SARS-CoV-2 RBD antigen for selection (see Methods); this resulted in enrichment of binders for all of the libraries except m396, as indicated by polyclonal phage ELISA (Fig. 3A).

**Figure 3.**
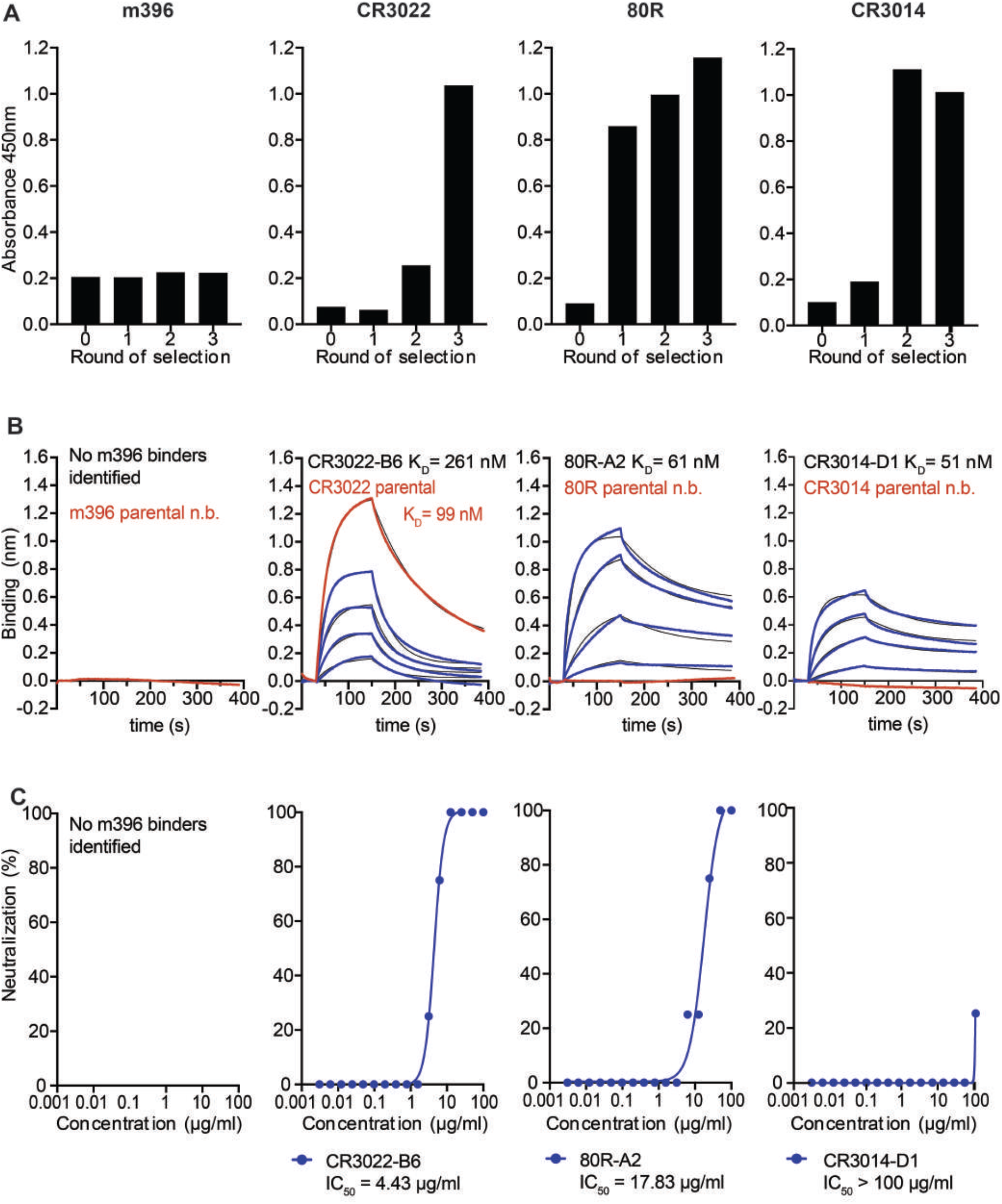
Selection of SARS-CoV-2 specific antibodies from light chain shuffled libraries of SARS-CoV-1 binders. (A) Enrichment of scFv antibody binders by phage display (polyclonal phage ELISA). Biolayer interferometry affinity measurements of soluble SARS-CoV-2 RBD (at 400 nM, 200 nM, 100 nM, 50 nM; highest concentration only shown for parental antibodies) binding to immobilized antibody (see Methods). (**C)** Neutralization of live SARS-CoV-2 virus in Vero E6 cells (IgG format).

In the case of the CR3022 selection, binders were dominated by a single clone (designated B6) after three rounds (no additional binders were identified when screening earlier selection rounds). The CR3022-B6 variant was expressed in an IgG format in CHO cells, and further characterized by biolayer-interferometry (BLI) and for neutralization in Vero E6 cells. These analyses revealed that CR3022-B6 IgG bound to soluble recombinant SARS-CoV-2 RBD with an equilibrium binding constant (K_D_) of 290 nM, lower than observed for the parental CR3022 antibody (99 nM) (Fig. 3B and Table 1). Intriguingly, and unlike parental CR3022, B6 was capable of neutralizing of live SARS-CoV-2 virus with an IC_50_ of 4.4 μg/ml (Fig. 3C). RBD mutagenesis was used to further characterize the CR3022-B6 epitope by targeting the CR3022-RBD interface through a K396S mutation designed to disrupt the interaction (Fig. 4D, surface C). While parental CR3022 binding was abolished through the mutation, CR3022-B6 fully maintained binding affinity (Supplementary Fig. 3B), indicating that the binding mode of the variant may have changed compared to the parental antibody. Next, CR3022-B6 was further improved through affinity maturation, by targeting all six CDR regions for diversification using Kunkel mutagenesis and off-rate selections on phage using biotinylated SARS-CoV-2 RBD (see Methods). This resulted in an affinity matured clone, designated CR3022-M, with moderately increased affinity (188 nM vs 290 nM for CR3022-B6) and considerably increased neutralization potential (0.35 μg/ml vs 4.4 μg/ml for CR3022-B6) (Supplementary Fig. 5).

**Figure 4.**
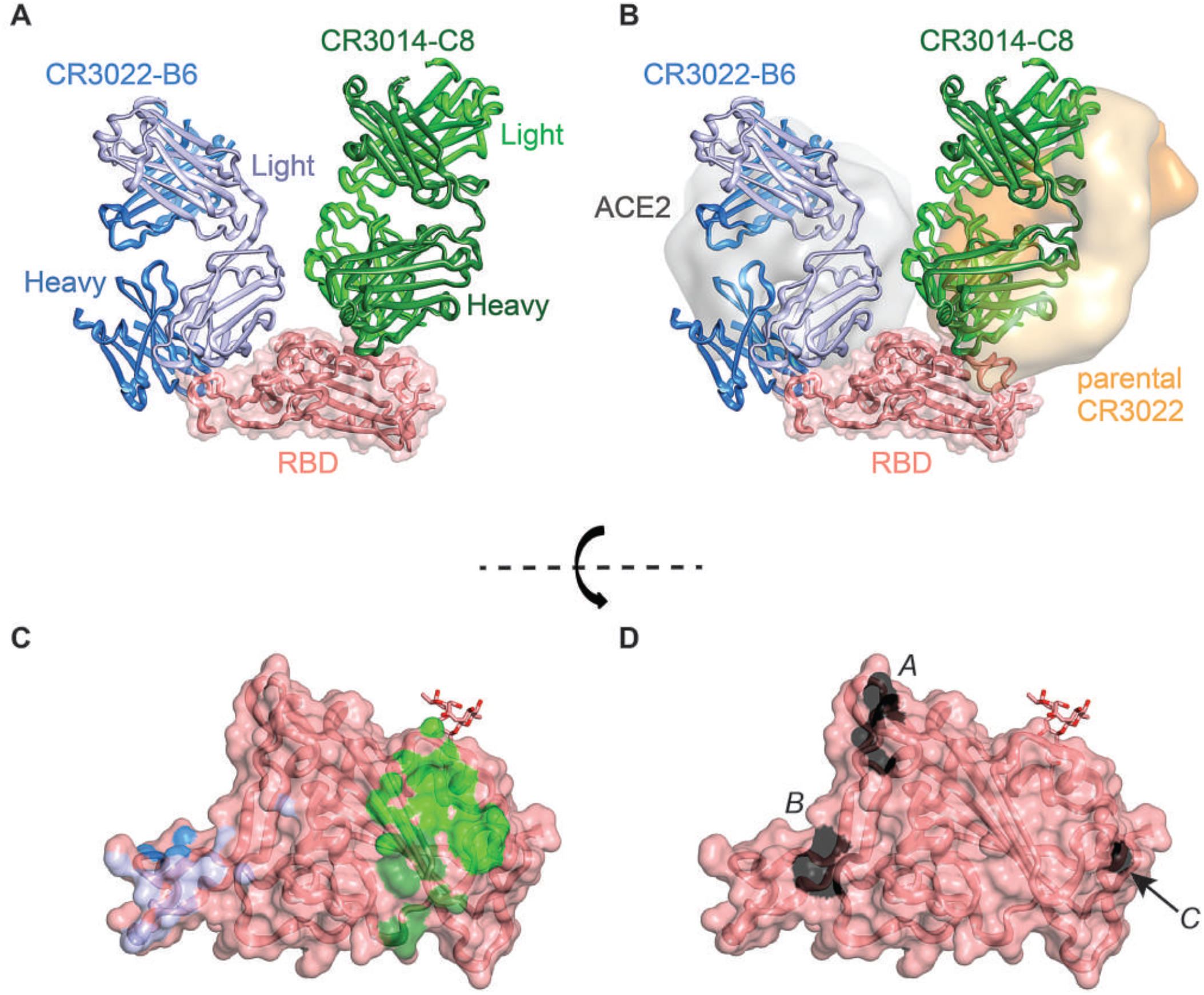
Crystal structure of CR3022-B6 and CR3014-C8 in complex with SARS-CoV-2 RBD. (**A)** CR3022-B6 (blue) and CR3014-C8 (green) Fabs bound to SARS-CoV-2 RBD (salmon); (**B)** CR3022-B6 binding to an RBD epitope overlapping with the ACE2 (grey surface) interface and different to the parental CR3022 (orange surfaces) epitope; CR3014-C8 binding to an epitope distant from the ACE2 interface; (B) Antibody contact surfaces on RBD for CR3022-B6 (blue) and CR3014-C8 (green). The majority of the RBD surface is buried by antibody V_L_ domains (light blue and light green), with more limited V_H_ interactions (dark blue and dark green); **(D)** RBD surface with residues targeted for epitope mapping in black; surface A (T500, N501 and Y505) ACE2 binding interface; surface B (L455 and F456) CR3022-B6 and CR3014-D1 interface (adjacent to the ACE2 binding site); surface C (K396) parental CR3022 binding interface.

In the case of the CR3014 selections, libraries became dominated by two clones (D1 and C8) after three rounds of selection. When converted into IgG (and unlike the parental CR3014 IgG, which did not detectably bind to SARS-CoV-2 RBD), both clones bound with mid-nanomolar affinity (KD of 51 nM and 61 nM respectively) (Fig. 3B and Table 1). Similar to the observation for the CR3022-B6/CR3022-parental pair, only CR3014-D1 (but not CR3014-C8) was able to detectably neutralize live virus (although weakly with an IC_50_ >100 μg/ml) (Fig. 3C).

In the case of the 80R selections, later rounds were dominated by a set of variants with closely related CDR sequences. A representative clone (80R-A2) was expressed in an IgG format and analyzed by BLI and neutralization: this revealed that the 80R-A2 variant had acquired the capacity to bind RBD and neutralize SARS-CoV-2 with a KD of 61 nM and an IC_50_ of 17.8 μg/ml (Fig. 3B-C and Table 1). Parental 80R has been shown to block the interaction of SARS-CoV-1 RBD with ACE2 ^14^. Similarly, 80R-A2 binding to SARS-CoV-2 RBD was disrupted through T500A/N501A/Y505A triple substitutions targeting the ACE2 binding site as described above (Fig. 4D surface A and Supplementary Fig. 3B).

### Structural characterization of light chain shuffled antibodies

In parallel to the affinity maturation of CR3022-B6, we crystallized this variant in a Fab format, both alone (1.7 Å) and as part of a ternary complex. The complex containing CR3022-B6 Fab, CR3014-C8 Fab and SARS-CoV-2 RBD diffracted to 2.8 Å (Fig. 4A). Although electron density for the CR3022-B6 Fab component in the ternary complex was weak (reflected in high average B factors - Supplementary Table 2), it was evident that the interaction was distant from the previously described binding site of parental CR3022 (Fig. 4B). This observation was further confirmed by targeting the CR3022-B6 RBD interface observed here with a double mutation in the RBD (L455A/F456A), which completely abolished binding of the variant (but not parental CR3022) (Fig. 4D surface B and Supplementary Fig. 3B). We next used cryo-electron microscopy (cryo-EM) to position the CR3022-B6 Fab onto the surface of SARS-CoV-2 spike trimer (Fig. 5A and Supplementary Table 3). In the absence of antibody, classification of the data indicated that the majority of spike trimers harbored two RBD domains in the down conformation, with the remaining RBD in the up position (Supplementary Fig. 6) ^23^. When incubated with an equimolar equivalent of CR3022-B6 Fab, ~77% of spike trimer particles were visualized with a Fab attached to RBD in an up conformation (Fig. 5A and Supplementary Fig. 6) and the remaining ~23% were in the single RBD up conformation. Comparison of the RBD-CR3022-B6 component of the crystallographic complex resulted in good agreement with the cryo-electron microscopy map, highlighting binding of CR3022-B6 in proximity to the RBD ACE2 binding site, but distant from the parental CR3022 binding site (Fig. 4B and Fig, 5A).

**Figure 5.**
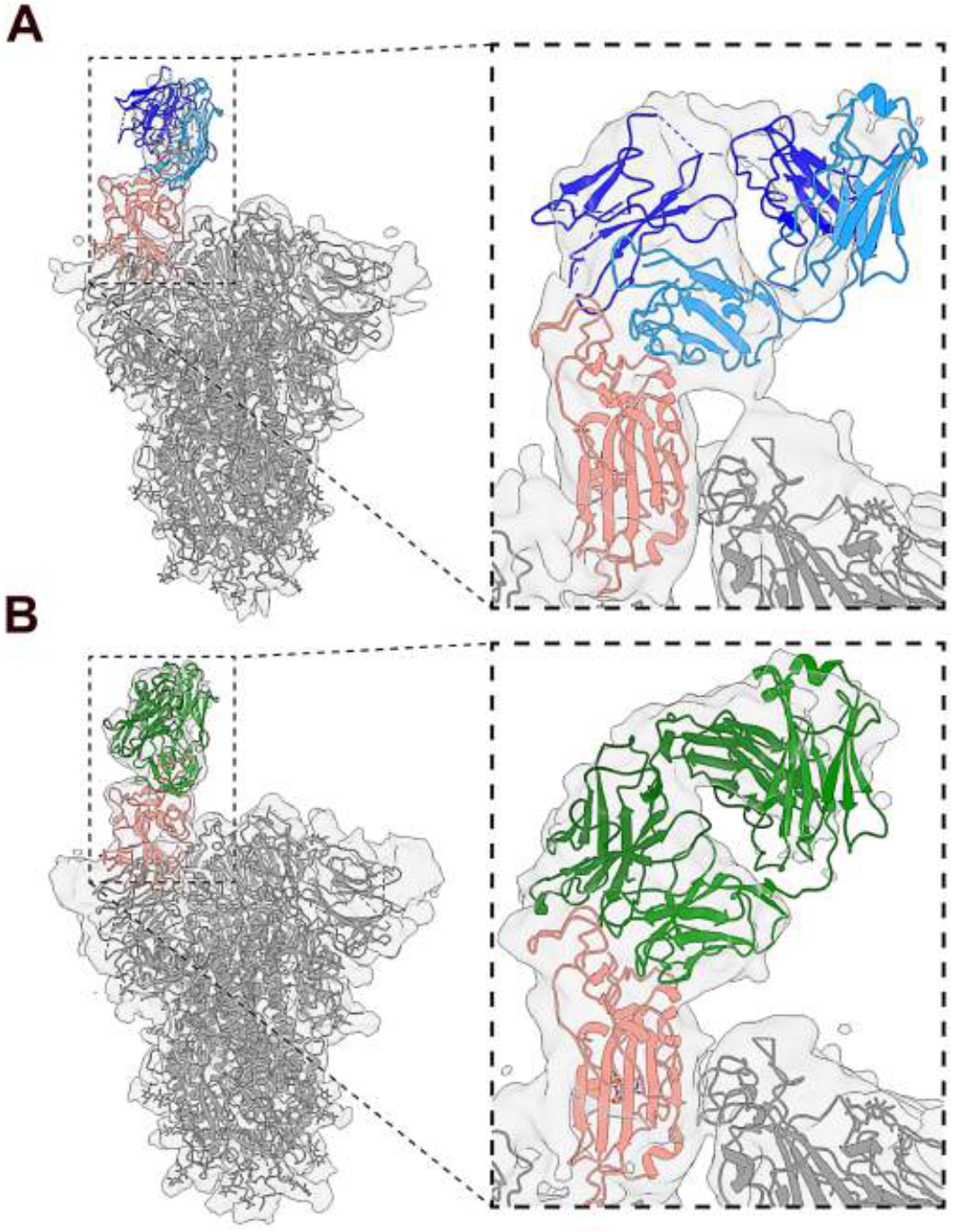
Cryo-electron microscopy of Fab-spike trimer complexes. (**A)** Binding of CR3022-B6 Fab (blue) to SARS-CoV-2 trimer (grey with RBD highlighted in salmon). A single Fab molecule bound to the RBD domain of a spike protomer in the ‘up’ conformation was resolved. (**B)** Binding of CR3014-D1 Fab (green) to SARS-CoV-2 trimer (grey with RBD highlighted in salmon). Both antibody Fab bound with similar stoichiometry (with two protomers in the ‘down’ and one in the ‘up’ conformation), and epitope specificty (overlapping with the ACE2 binding interface).

In contrast to CR3022-B6, electron density of the CR3014-C8 Fab component was well defined in the crystal structure of the ternary complex, highlighting an epitope distant from the ACE2 binding site (Fig. 4A). Indeed, the epitope of CR3014-C8 closely resembled that of the CR3022 antibody (Fig. 4B). Further structural information for a second, but neutralizing, CR3014 variant (CR3014-D1) was obtained by cryo-EM. When incubated with an equimolar equivalent of CR3014-D1 Fab, ~45% of spike particles were visualized with a single Fab attached to an up-conformation RBD domain (Fig. 5B and Supplementary Fig. 6), with the remaining particles in the single RBD up conformation. Superposition of the CR3022-B6 and CR3014-D1 cryo-EM spike-Fab structures revealed that both Fabs bound to highly similar and overlapping RBD epitopes (Supplementary Fig. 7). These observations were in agreement with mutagenesis experiments, with both CR3022-B6 and CR3014-D1 binding abolished by mutations (L455A/F456A) adjacent to the RBD ACE2 binding site (Fig. 4D, surface B), while CR3014-C8 binding was abolished by a mutation centered on the RBD CR3022 binding site (K396S) (Fig. 4D, surface C, and Supplementary Fig. 3B).

## DISCUSSION

Here we describe a rapid and straightforward *in vitro* strategy for the generation of antibodies that potently bind and neutralize SARS-CoV-2. Utilizing two complementary strategies, site-directed mutagenesis and light chain shuffling, we introduced diversity into the variable domain regions of four well characterized monoclonal antibodies that had been developed after the 2003 SARS outbreak (Fig. 1A-B).

From the site-directed mutagenesis libraries we identified variants of antibodies m396, CR3022 and 80R that bound to SARS-CoV-2 RBD by soluble ELISA (no binders were obtained for CR3104 selections). A variant of m396, m396-B10 was further characterised by biolayer interferometry. These experiments revealed that while the parental m396 antibody displayed no detectable binding, m396-B10 bound to SARS-CoV-2 RBD with single digit nanomolar monovalent affinity (Fig. 2; compared to 20 nM for the m396-parental/SARS-CoV-1 RBD interaction ^21^). Neutralization of live SARS-CoV-2 virus in Vero E6 cells confirmed potent neutralization, with an IC_50_ of 160 ng/ml. Neutralization was also observed in SARS-CoV-2 and SARS-CoV-1 pseudoparticle assays indicating potent cross-specificity of m396-B10.

In contrast, no detectable increase of equilibrium binding affinity was observed for variants of CR3022, although two of the analysed variants displayed increased *kinetic* association constants for SARS-CoV-2 binding compared to the parental CR3022 antibody. However, none of the CR3022 variants generated by site-directed mutagenesis displayed detectable viral neutralization (Fig. 2C).

From the kappa light chain shuffled libraries we identified variants of antibodies CR3022, 80R and CR3014 that bound to SARS-CoV-2 RBD (no binders were obtained for m396 selections, presumably due to the presence of lambda light chain in the parental antibody). The CR3022 selections were dominated by a single clone, CR3022-B6, which bound SARS-CoV-2 with reduced affinity compared to the parental CR03022 antibody (290 nM compared to 99 nM) (Fig. 3B and Table 1). Intriguingly, and unlike wild-type CR3022, CR3022-B6 effectively neutralized SARS-CoV-2 live virus with an IC_50_ of 4.4 μg/ml. This apparent discrepancy between affinity and neutralization potential in otherwise closely related variants was further confirmed through epitope mapping. Mutation of the SARS-CoV-2 RBD CR3022 binding site abolished binding of recombinant parental CR3022, but not of CR3022-B6 (Supplementary Fig. 3B). In contrast, CR3022-B6 binding was abolished by mutation of the RBD adjacent to the ACE2 binding site (L455A/F456A, Fig. 4D surface B), which did not affect binding of CR3022 wild type (Supplementary Fig. 3B). The suggestion that CR3022-B6 binds a different epitope to its parent was confirmed by a crystal structure of CR3022-B6 in complex with SARS-CoV-2 RBD (along with CR3014-C8 in a ternary complex), and through cryo-electron microscopy analysis of CR3022-B6 in complex with SARS-CoV-2 trimeric spike. Both structural analyses confirmed binding of CR3022-B6 in proximity to the RBD ACE2 interaction surface, and distant to the original CR3022 binding site, providing a rationale for its observed neutralization activity.

The observation that a human antibody, in its wild-type and matured light chain shuffled form, can bind to two completely different epitopes is intriguing. This observation was not limited to CR3022, with similarly distinct epitopes observed among two variants of the CR3014 antibody (CR3014-C8 and CR3014-D1). Despite the absence of structural information for the parental CR3014 antibody, mutagenesis, crystallography and cryo-electron microscopy clearly highlight the considerable difference in epitope specificity for these two light chains shuffled clones. The difference in specificity also correlates well with neutralization potential: clones binding to a surface proximal to the ACE2 binding site (CR3022-B6 and CR3014-D1) display neutralization activity against live SARS-CoV-2, while clones binding distant to the ACE2 binding site are not detectably neutralizing (parental CR3022 and CR3014-C8).

How are antibodies with identical heavy chain, but different light chains, capable of binding to two completely different epitopes? Further inspection of the CR3022-B6 and CR3014-C8 interactions with SARS-CoV-2 RBD, and the relative contributions of their heavy and light chains to the interaction, provide an intriguing insight into this question: while the majority of the interface (and binding energy) in most antibody-antigen interactions is dominated by variable heavy (VH) domains, this is not the case in the variant structures reported here (Fig. 4C). In the case of CR3022-B6 a total of 480 Å^2^ of buried surface is observed for V_L_, with only 230 Å^2^ observed for V_H_. (relative contact surfaces shaded blue in Fig. 4C). This is marked contrast to the parental CR3022 antibody where the interaction is dominated by heavy chain contacts (V_H_ : 592 Å^2^ and V_L_ : 415 Å^2^). Similar ratios were observed for the CR3014-C8 interaction with SARS-CoV-2 (V_H_ : 240 Å^2^ and V_L_ : 460 Å^2^, Fig 4C, green shading). In the case of CR3022-B6, the increase of V_L_ contact surface is accompanied by a considerable reduction in the length of CDR L1, which is elongated in the parental CR3022 structure, resulting in a more shallow and extended interface and allowing for additional interactions (Supplementary Figure 2B).

Taken together, our results demonstrate that high affinity antibodies against a variant coronavirus can be generated through maturation of previously reported antibodies. Several of the selected antibodies potently neutralized live SARS-CoV-2 virus assays with half-maximal inhibitory concentrations (IC_50_) in the nanomolar range, well within the therapeutic range ^23^.

While the use of site-directed mutagenesis libraries resulted in the selection of potent variants with conserved epitope specificity, the use of light chain shuffling also resulted in the generation of antibodies with completely new specificities. The discovery of such dual specificity antibody pairs with identical heavy chains, but different light chains, is intriguing and may enable the generation of bi-specific reagents with improved resistance against mutational escape ^24^.

The observation that a limited number of CDR mutations can endow nanomolar affinity binding and potent neutralization onto antibodies originally raised against a different variant coronavirus (SARS-CoV-1), also raises important implications for natural immunity and vaccine design. While the potential of antibody maturation against variant antigens has been demonstrated using haptens ^25^ and viral model antigens ^26^, insights into the mutational plasticity of coronavirus antibodies had remained unclear. We conclude that *in vitro* maturation provides a rapid pathway for the identification of potent antibody reagents against emerging viruses.

## METHODS

### Generation of site-directed mutagenesis antibody libraries

m396, CR3022, CR3014, and 80R scFv were gene synthesized (Genscript) and cloned into the pHEN1 phagemid vector. Site-directed mutagenesis was carried out by Kunkel mutagenesis ^16^. In brief, phagemid vectors were transformed into *E. coli* CJ236, single colonies grown in 2xYT media supplemented with 100 μg/mL ampicillin, 10 μg/mL chloramphenicol and 2% glucose until reaching an OD_600nm_ of 0.4. Bacteria were then infected with KM13 helper phage and grown overnight at 30°C in 2xYT media supplemented with 100 μg/mL ampicillin, 10 μg/mL chloramphenicol, 50 μg/mL kanamycin and 0.25 μg/mL uridine. Phage particles were purified from the culture media using PEG/NaCl and uridine containing single-stranded DNA (dU-ssDNA) extracted using a QIAprep spin M13 kit (Qiagen). Mutagenesis was carried out by annealing degenerated oligonucleotides to the dU-ssDNA, followed by synthesis of the covalently closed circular DNA (cccDNA) with T7 DNA polymerase and T4 ligase (NEB). Finally, the cccDNA was transformed into electro-competent *E. coli* TG1 and bacteria titrated to determine library sizes. Bacteria were harvested from the agar plates, grown in 2xYT media supplemented with 100 μg/mL ampicillin, and 2% glucose until reaching an OD_600nm_ of 0.4. At this point, bacteria were infected with KM13 helper phage and grown overnight at 30°C in 2xYT media supplemented with 100 μg/mL ampicillin, and 50 μg/mL kanamycin. Phage antibody libraries were purified from culture media using PEG/NaCl and stored at 4°C.

### Generation of light chain shuffled antibody libraries

DNA encoding SARS-CoV-1 V_H_ regions was amplified by PCR (using Q5 polymerase NEB). J segments were modified as required to match the following protein sequence (GTLVTVSS). Region encoding V kappa library regions were amplified from a pHEN1 scFv library, comprising the end of the V_H_ J segment, glycine-serine linker and V_L_ regions. The resulting light chain shuffled library was generated by splice-overlapping extension PCR and cloned into pHEN1 in a scFv format using NcoI and NotI restriction sites. DNA was transformed into electro-competent TG1, and phage produced and purified as above.

### Phage display selections

For phage display selection, we biotinylated SARS-CoV-2 RBD using a terminal AviTag and BirA biotin ligase (Avidity) according to the manufacturer’s protocol. Phage display selections were carried out by alternating between capture of the antigen on neutravidin coated wells on Maxisorp plates (Nunc) and streptavidin magnetic beads (Invitrogen) ^27^. For Maxisorp plate selection, neutravidin was coated overnight at 50 μg/mL in carbonate coating buffer, biotinylated RBD captured, and blocked in PBS supplemented with 0.1% Tween-20 and 4% skim milk (MPBST). 1 × 10^12^ phage were blocked in MPBST, added to the wells containing antigen and incubated for 1 h. The wells were washed with 1xPBST, 1xPBS. Phage were eluted with 100 μg/mL trypsin for 1 h, then used to infect TG1 bacteria at an OD_600nm_ of 0.4. Infected TG1 were plated onto 2xYT agar plates supplemented with 100 μg/mL ampicillin and 2% glucose. For streptavidin beads selection, phages were blocked as described above and incubated with biotinylated RBD. 30 μl of streptavidin magnetic beads (Invitrogen) were blocked in PBST supplemented with 4% BSA (Sigma), then incubated for 15 min with the phage/antigen mix. Magnetic beads were washed with PBST and PBS and phage eluted as described above. For site-directed mutagenesis libraries, we used 100 nM, 50 nM, 5 nM and 0.5 nM of biotinylated RBD for selection rounds 1 to 4, and 100 nM, 50 nM, 25 nM and 10 nM for light chain shuffled libraries. Phage titres used for selection were reduced to 1 × 10^11^ for rounds 2 and 3 and 1 × 10^10^ for round 4.

Affinity maturation was carried out using off-rate selections and streptavidin magnetic beads. Selections were performed essentially as previously described^28^, with the following adjustments: phage were incubated with the biotinylated RBD for 1h, excess unbiotinylated RBD was added (100x and 350x for rounds 2 and 3) and further incubated for 2/8 h for rounds 2/3 before capture on magnetic streptavidin beads.

### Polyclonal phage and monoclonal soluble ELISA

For polyclonal ELISA, Maxisorp plates were coated with neutravidin overnight and 100 nM of biotinylated RBD was subsequently captured. 1 × 10^9^ purified phage were blocked in MPBST and incubated in each well for 1h. Plates were washed with PBST, incubated with HRP-conjugated anti-M13 antibody (GE Healthcare) for 1h and washed again. The plate was finally incubated with TMB substrate (Perkin Elmer), the reaction quenched with HCl and the plate read at Abs_450nm_ (ClarioStar – BMG Labtech). For monoclonal soluble ELISA, individual colonies from the selection titration plates were inoculated in 96 well plates and incubated at 37°C overnight. The bacteria were re-inoculated the following day at 1:50 and incubated at 37°C for 4h. The plates were then spun down, the culture media discarded, bacteria resuspended in 2xYT supplemented with 100 μg/ml ampicillin and 1 mM IPTG and incubated overnight at 30°C. For ELISA, Maxisorp plates were coated with neutravidin overnight and 100 nM of biotinylated RBD subsequently captured. The plates were then incubated with 50 μl of culture media, clarified by centrifugation, for 1h and then washed with PBST. The plates were subsequently incubated with HRP-conjugated chicken anti c-myc antibody (ICL Lab) for 1h and washed again. The plate was finally incubated with TMB substrate (Perkin Elmer), the reaction quenched HCl and the plate read at Abs_450nm_ (ClarioStar – BMG Labtech).

### Monoclonal antibody production and purification

DNA encoding antibody variable domains was amplified by PCR from the pHEN1 phage display vector and cloned into a human IgG1 expression vector based on pCEP4 (Invitrogen). After validation of the cloning by Sanger sequencing, the plasmids were transfected into ExpiCHO cells (Thermo Scientific) according to the manufacturer’s protocol (1 μg DNA/ml of cells; 2:1 ratio of heavy chain to light chain) and following the max titer protocol. After 14 days, cell culture media were clarified by centrifugation and the IgG captured using Protein G resin (Genscript). IgG were eluted from the resin using 100 mM glycine pH 3.0, eluate was dialyzed against PBS the purity assessed by SDS-PAGE. For Fab production, DNA encoding V_H_ and V_L_ regions was cloned into a pCEP4 based vector encoding a C-terminal His tag. Production was carried out in ExpiCHO cell as above. After 14 days, cell culture media were clarified by centrifugation, dialyzed against PBS and Fab protein captured using Talon Resin (Thermo Scientific). Fab protein was eluted with 150 mM imidazole in PBS, dialyzed with PBS and the purity assessed by visualization on SDS-PAGE. In the case of m396-B10, Fab was generated through proteolytic cleavage of IgG using papain and purified using protein A affinity chromatography.

### Affinity measurements using biolayer interferometry (BLI)

Purified monoclonal antibodies (Fab/IgG) were buffer exchanged into PBS using equilibrated ZebaSpin columns (Thermo Fisher Scientific). The protein concentration was determined and the antibodies biotinylated by incubating for 30 min at room temperature with EZ-Link NHS-PEG4-Biotinylation reagent (Thermo Fisher Scientific) at a 10:1 biotin-to-protein ratio. Free biotin was removed from the samples by repeating the buffer exchange step in a second ZebaSpin column equilibrated with PBS. Affinity of interactions between biotinylated antibodies and purified soluble RBD proteins were measured Biolayer Interferometry (BLItz, ForteBio). Streptavidin biosensors were rehydrated in PBS containing 0.1% w/v BSA for 1 h at room temperature. Biotinylated antibody was loaded onto the sensors “on-line” using an advanced kinetics protocol, and global fits were obtained for the binding kinetics by running associations and dissociations of RBD proteins at a suitable range of molar concentrations (2-fold serial dilution ranging from 800 nM to 50 nM). The global dissociation constant (K_D_) for each 1:1 binding interaction was determined using the BlitzPro 1.2.1.3 software. Human IgG1 was used for all measurements, except for 80R-A2, CR3014-D1, CR3014-C8 for which Fab was used. For ACE2 competition assay, biotinylated ACE2-Fc was loaded onto the streptavidin sensors on-line, and the binding kinetics determined using either 500 nM of soluble RBD, or 500 nM of soluble RBD pre-incubated with 1 μM of IgG for 5min, using advanced kinetics protocol.

### Antigen production and purification

DNA encoding SARS-Cov-2 RBD (residues 319-541) was gene synthesized (Genscript) and cloned into pCEP4 mammalian expression vector with a N-terminal IgG leader sequence and C-terminal Avitag and His tag. The plasmid was transfected into Expi293 cells (Thermo Scientific) according to the manufacturer’s protocol and the protein expressed for 7 days at 37°C, 5% CO_2_. The cell culture was clarified by centrifugation, dialyzed with PBS and the protein captured with Talon resin. The RBD was eluted with 150 mM imidazole in PBS, dialyzed with PBS and the purity assessed by visualization on SDS-PAGE. The plasmid encoding the spike protein with C-terminal trimerization domain and His tag was a gift from the Krammer lab (BEI Resources). The plasmid was transfected into Expi293 cells and protein expressed for 3 days at 37°C, 5% CO_2_. The protein was purified using the His tag as for the RBD purification. The protein was further purified on a Superose 6 gel filtration column (GE Healthcare) using an AKTA Pure FPLC instrument (GE Healthcare) to isolate the trimeric protein and remove S2 pre-fusion protein.

### SARS-CoV-2 neutralization assays

Serial 2-fold dilutions of test monoclonal antibody were prepared in 96-well plates in octuplicate. The serial dilutions were incubated for 1 hour at 37°C with an equal volume of SARS-CoV-2 isolate containing 200 TCID_50_ (infectious dose). A Vero E6 suspension containing 2 × 10^4^ cells was added to each well, and plates were incubated at 37°C (5% CO_2_). After 3 days, the plates were observed for cytopathic effect (CPE) and IC_50_ values were calculated from four parameter dose-response curves (GraphPad Prism). All dilution steps of antibody, virus, and cells were performed in culture media containing MEM, 2% fetal bovine serum, and 1x penicillin-streptomycin-glutamine.

### SARS-CoV pseudovirus neutralisation assays

Cells were cultured at 37°C and 5% CO_2_ in growth medium containing high glucose Dulbecco’s Modified Eagle Medium (ThermoFisher Scientific) supplemented with 10% (v/v) heat inactivated fetal bovine serum (Life Technologies; ThermoFisher Scientific). Retroviral SARS-CoV-1 and 2 pseudo-particles (SARS-2pp) were generated by co-transfecting expression plasmids containing SARS-CoV-1 or SARS-CoV-1 spike which were kindly provided by Prof Gary Whitaker and Dr Markus Hoffmann^r^, respectively, and the MLV gag/pol and luciferase vectors which were kindly provided by Prof. Francois-Loic Cosset, in CD81KO 293T cells, which were kindly provided by Dr Joe Grove^30^, using mammalian Calphos transfection kit (Takara Bio). Culture supernatants containing SARS-2pp were harvested 48 hours post transfection and clarified of cellular debris by centrifugation at 500xg for 10 minutes. SARS-2pp were concentrated 10-fold using 100,000 MWCO Vivaspin centrifugal concentrators (Sartorius) by centrifugation at 2000xg and stored at –80°C. For neutralisation assays, the infectivity of SARSpp were diluted in media to 1000 – 5000-fold more infectious than negative background (based on pseudoparticles lacking SARS-CoV Spike). Diluted pseudoparticles were incubated for one hour with monoclonal antibodies, followed by the addition of polybrene at a final concentration of 4μg/mL (Sigma-Aldrich), prior to addition to 293T-ACE2 over-expressed calls, which were kindly provided by A/Prof Jesse Bloom. 293T-ACE2 cells were seeded 24 hours earlier at 1.5 × 10^4^ cells per well in 96-well white flat bottom plates (Sigma-Aldrich). Cells were spinoculated at 800xg for two hours and incubated for two hours at 37°C, prior to media change. After 72 hours, the cells were lysed with a lysis buffer (Promega) and Bright Glo reagent (Promega) was added at a 1:1 ratio. Luminescence (RLU) was measured using CLARIOstar microplate reader (BMG Labtech). Neutralisation assays were performed in triplicates and outliers were excluded using the modified z-score method. Percentage neutralisation of SARSpp was calculated as (1 – RLU_treatment_/RLU_no treatment_) × 100. The 50% inhibitory concentration (IC50) titre was calculated using non-linear regression model (GraphPad Prism).

### X-ray crystallography

Gel-filtration chromatography purified SARS-CoV-2 RBD (residues 333-528), and light chain shuffled Fabs CR3022-B6 and CR3014-C8 (in 25 mM Tris pH 8.0, 200 mM NaCl) were combined in a 1:1:1 molar ratio at a concentration of ~4 mg/mL, from which equal volumes (2 μL) were combined with well solution (100 mM Tris pH 8.0, 200 mM NaCl and 20% PEG3350) in a hanging drop format. After several weeks, crystals of sword-like morphology appeared growing out of precipitate. Due to their small size crystals were harvested without a cryoprotection regime and plunge-frozen in liquid nitrogen. Large crystals of CR3022-B6 Fab alone were grown by combining equal volumes (2 μL) of protein (9.6 mg/mL) with well solution comprising 200 mM sodium citrate (pH 6.65) and 24% PEG3350. Crystals of m396-B10 were grown by combining equal volumes of protein (5.95 mg/ml) with well solution comprising 200 mM NaCl, 100 mM BisTris (pH 5.95) and 25% (v/v) PEG 3350. Cryoprotection for CR3022-B6 crystals was achieved by briefly (5-10s) swimming crystals in well solution supplemented with glycerol (to ~25% v/v) prior to looping and snap freezing. Diffraction data were collected at the Australian Synchrotron on beamline MX2 using a Dectris Eiger X16M detector. In both cases a 360° sweep of data were deconvoluted into 3600 × 0.1° oscillation images which were indexed and integrated by XDS ^29^. Space groups were determined with Pointless ^30^ and scaling and merging performed with Aimless ^31^, both components of CCP4 ^32^.

Structures were determined by molecular replacement using Phaser ^33^. The search model for the CR3022-B6 structure was the CR3022 Fab component of PDB entry 6w41 ^15^, split into variable domain (V_H_ + V_L_) and constant domain (CH1 + CL) pairings. For the double-Fab 1:1:1 complex (RBD + CR3014-C8 + CR3022-B6), cell content analysis suggested that, should all components be present in the expected stoichiometric ratio, the solvent content would be 50%. The search model for the RBD was also derived from PDB entry 6w41. The search model for the Fab components were the same V_H_ + V_L_ and CH1 + CL pairings as used for the CR3022-B6 structure alone. Molecular replacement was able to place two Fabs without clashing on the surface of a single RBD, although the second Fab returned a much smaller log likelihood gain than the first. This was reflected in electron density for one Fab being unambiguous and well resolved (clearly CR3014-C8), whilst the other was very weak, suggesting either incomplete occupancy or conformational/positional motion of this Fab within the lattice. CR3022-B6 was modeled into this second Fab position. Interestingly, the CR3014-C8 Fab bound RBD where wild-type CR3022 would have been expected to bind, whilst CR3022-B6 bound to a surface of RBD consistent with neutralization assay data suggesting it was in fact neutralizing (unlike parental CR3022). Interestingly, the bulk of buried surface for both interactions was dominated by the light chains (Supplementary Table 2), which for these Fabs are very similar, raising the issue of whether the CR3022-B6 Fab was in fact correctly modeled (the V_H_ region, in particular, was very poorly resolved). That CR3022-B6 does bind this second position was confirmed by a combination of; fo-fc difference maps suggesting CR3022-B6 was a better fit, molecular replacement yielding stronger solutions with CR3022-B6 placement relative to CR3104-C8 placement, mutagenesis of the RBD epitope eliminating CR3022-B6 binding, and cryo-EM returning a spike + CR3022-B6 model consistent with CR3022-B6 binding this epitope. A short branched chain carbohydrate was clearly present attached to Asn343 of the RBD, and has been modeled as two N-acetyl glucosamine (NAG) sugars connected by a beta(1-4) glycosidic bond, with a fucose (FUC) residue attached to the N-linked NAG via a beta(1-6) glycosidic linkage. For the m396-B10 structure, the search model was the Fab component of PDB ID 2g75. Diffraction data and model refinement statistics are shown in Supplementary Table 2.

### Cryo-electron microscopy

For sample preparation, either spike trimer alone or 1:1 Fab:monomer (molar ratio - for the CR3022-B6 or CR3014-D1 datasets) was incubated at room temperature for 1 h before applying to holey gold grids and freezing. 3.5 μl of each sample was applied to 1.2/1.3 Ultrfoil Au grids (Quantifoil) which had been glow-discharged for 1 min at 19 mA. Plunge freezing was performed using a Vitrobot Mark IV (ThermoFisher) with 0 blot force, 4 s blot time and 100% humidity at 22 °C. For data collection, grids were transferred to a Talos Arctica Electron Microscope (ThermoFisher) operating at 200 kV equipped with a FalconIII direct detector. Movies were recorded using EPU with a calibrated pixel size of 0.986, a total dose of 40 electrons spread over 29 frames and a total exposure time of 60 s. For data processing, motion correction, CTF estimation ^34^ and blob particle picking were performed in cryoSPARC ^35^. Extracted particles were subjected to multiple rounds of 2D classification and ab initio reconstruction in cryoSPARC before their locations were exported to Relion 3.0 ^36^. Motion correction and CTF estimation was then implemented in Relion 3.0 and particles were reextracted and again subjected to 2D classification before 3D auto-refinement and Bayesian polishing. 3D classification was then used to sort the particles based on whether density (attributed to Fab) was present above on of the RBDs. The final Fab bound trimer particles were then imported back to cryoSPARC for NU-3D refinement. Supplementary Fig. 6 provides a flowchart to describe this workflow along with FSC curves. Supplementary Table 3 provides a summary of the data collection and refinement statistics.

## ACKNOWLEDGEMENTS

We thank the UTS Biologics Innovation Facility for help on SARS-CoV-2 spike protein production, and Pamela J. Bjorkman and Christopher Barnes (Caltech) for advice and help with early structural studies. This work was supported by the Medical Research Future Fund 2020 Antiviral Development Call (2001739); by Program Grants 1016953 and 1113904; Project Grant 1108800; Fellowships 585490, 1157744, 190774, 1176351, and 1081858 from the National Health and Medical Research Council; Discovery Grant 160104915; DECRA Fellowship DE190100985 from the Australian Research Council.

## AUTHOR CONTRIBUTIONS

R.R., O.M., P.S., and D.C. designed the project; R.R. performed light chain shuffling, selection and maturation; O.M. performed site-directed mutagenesis and selection; D.B.L. performed X-ray crystallography; M.S. and A.G.S. performed cryo-EM; G.W., A.A., R.A.B. and W.D.R. performed neutralization experiments; O.M., R.R., J.J., S.U. and J.Y.H. performed protein production and purification; H.L. and P.S. performed affinity measurements; R.R., D.B.L. and D.C. wrote the manuscript with input from all other authors.

---

**Supplementary Figure 1.**
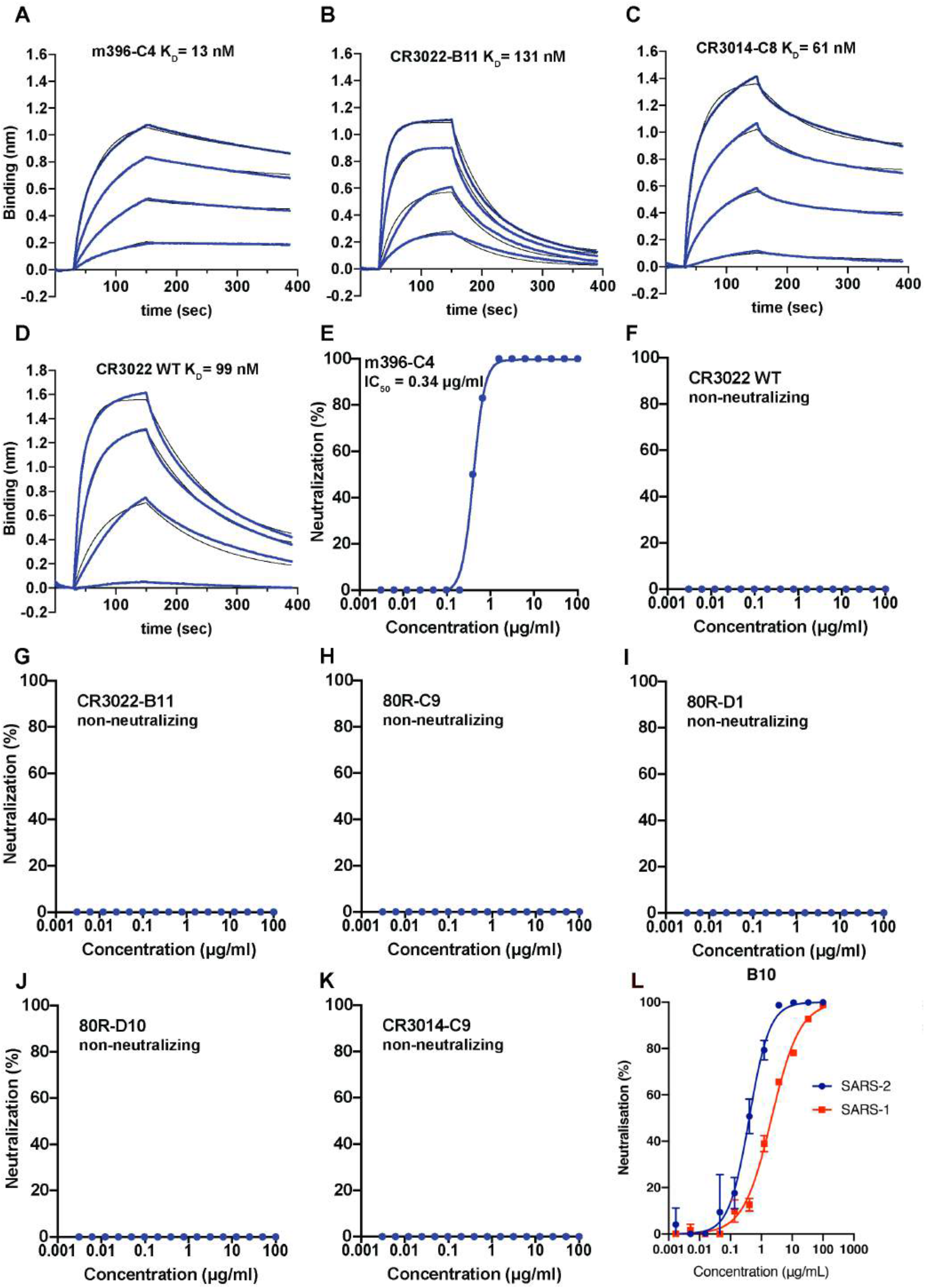
Affinity and neutralization measurements of matured antibodies. Affinity measurements of the monoclonal antibodies binding to SARS-CoV-2 RBD by biolayer interferometry. Biotinylated antibody (see Methods) of **(A)** m396-C4; **(B)** CR3022-B11; **(C)** CR3014-C8; **(D)** parental CR3022 was immobilized onto streptavidin sensors and incubated with different concentrations of SARS-CoV-2 RBD (400 nM to 50 nM in 2-fold dilutions) (see Table 1 for affinities); **(E-K)** Neutralization of SARS-CoV-2. Serial dilutions of IgG and SARS-CoV-2 were pre-incubated for 1h at 37°C and incubated with Human Vero E6 cells for 3 days; (L) Neutralization of SARS-CoV-1 and SARS-CoV-2 pseudoparticles by m396-B10 (IC_50_2.2 and 0.3 μg/ml).

**Supplementary Figure 2.**
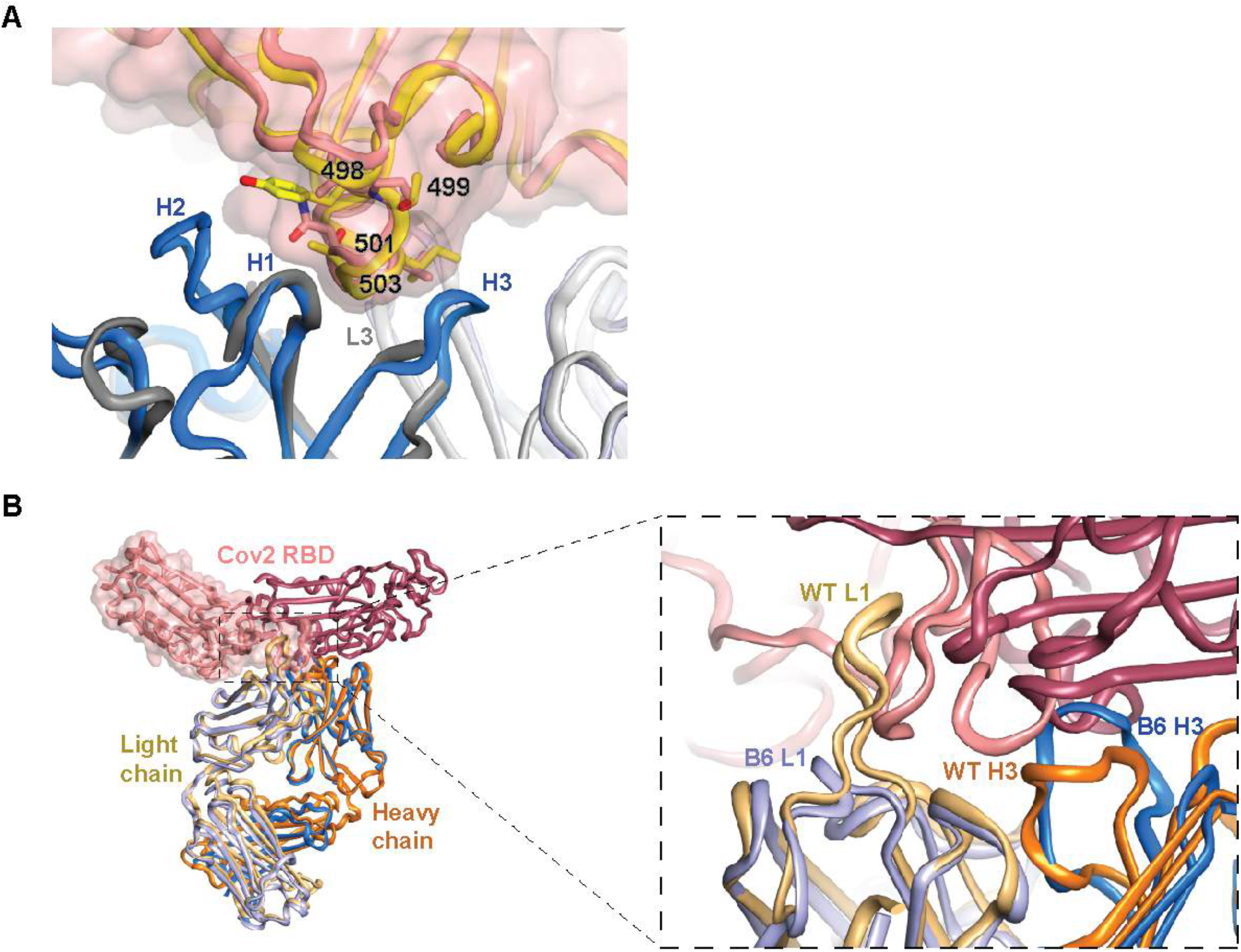
m396 and CR3022 interfaces. **(A)** Crystal structure of m396-B10 Fab in isolation (heavy and light chains colored dark and light grey) superposed with the Fab component of the complex between m396 (heavy and light chains colored dark and light blue) and SARS-CoV-1 RBD (yellow cartoon)(SARS-CoV-2 RBD superposed in salmon cartoon and transparent surface). Disorder within the electron density prevents tracing the fold of m396-B10 CDRs H3 and H2 regions. Numbered residues and accompanying sticks indicate positions within the RBD that are divergent between SARS-CoV-1 and SARS-CoV-2. **(B)** Superposition of the RBD-Fab components of parental CR3022 (heavy and light chains colored orange and light orange) bound to SARS-CoV-2 RBD (burgundy cartoon), and CR3022-B6 (heavy and light chains colored blue and light blue) bound to SARS-CoV-2 RBD (salmon cartoon and transparent surface). The right-hand panel showing a close up view of the considerably different RBD surfaces targeted by the two antibodies (of particular note are the different length of CDR L1 and the different conformation of CDR H3, both accommodating different RBD features).

**Supplementary Figure 3.**
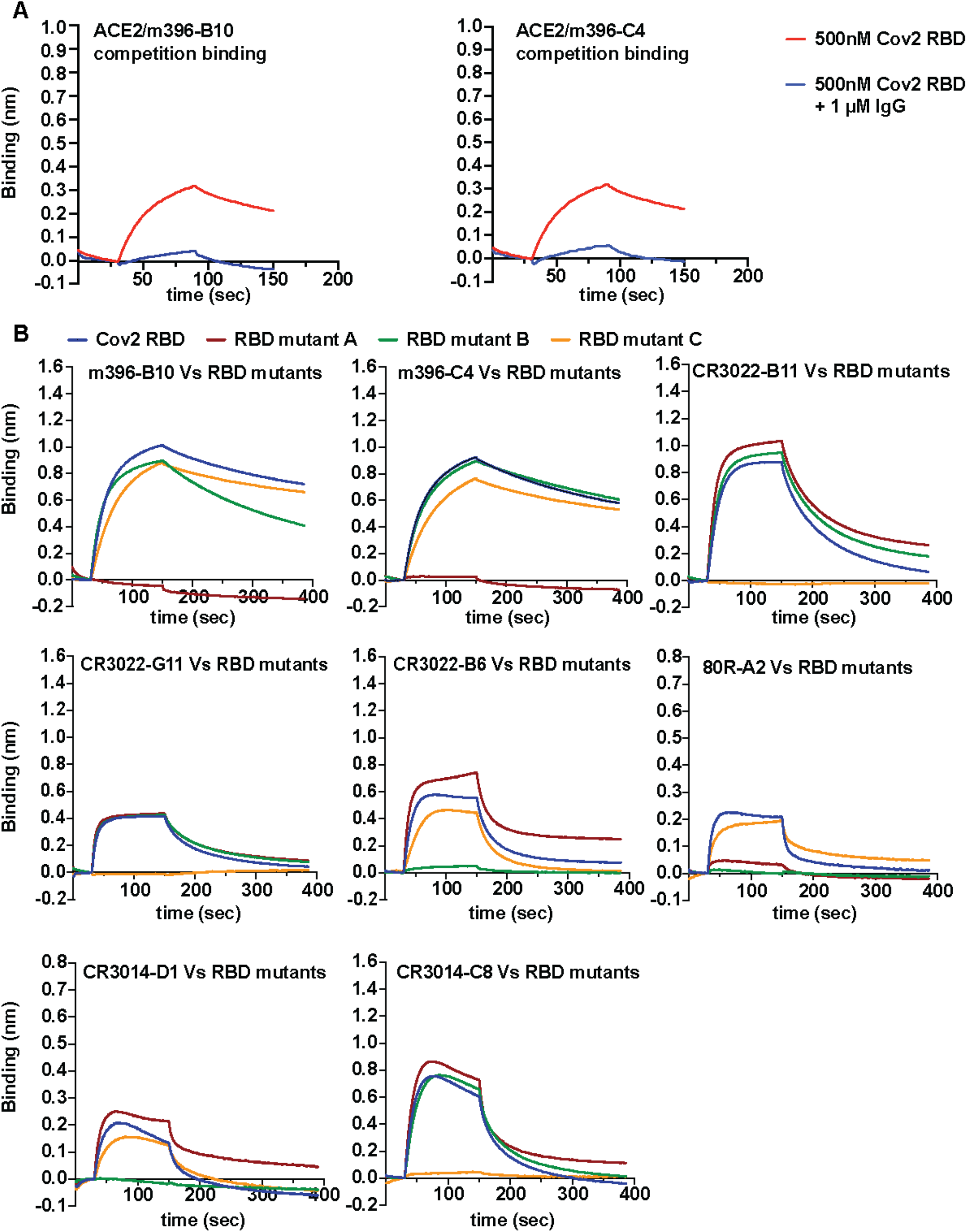
Epitope mapping. **(A)** ACE2 competition was investigated by BLI: biotinylated ACE2-Fc was immobilized onto streptavidin sensors and incubated with either SARS-CoV-2 RBD at 500 nM or with SARS-CoV-2 RBD at 500 nM pre-incubated with IgG at 1 μM; **(B)** Biotinylated IgG immobilized onto streptavidin sensors and incubated with SARS-CoV-2 RBD wild-type, mutant A (T500A, N501A and Y505A), mutant B (L455A and F456A) or mutant C (K396S) at 400 nM.

**Supplementary Figure 4.**
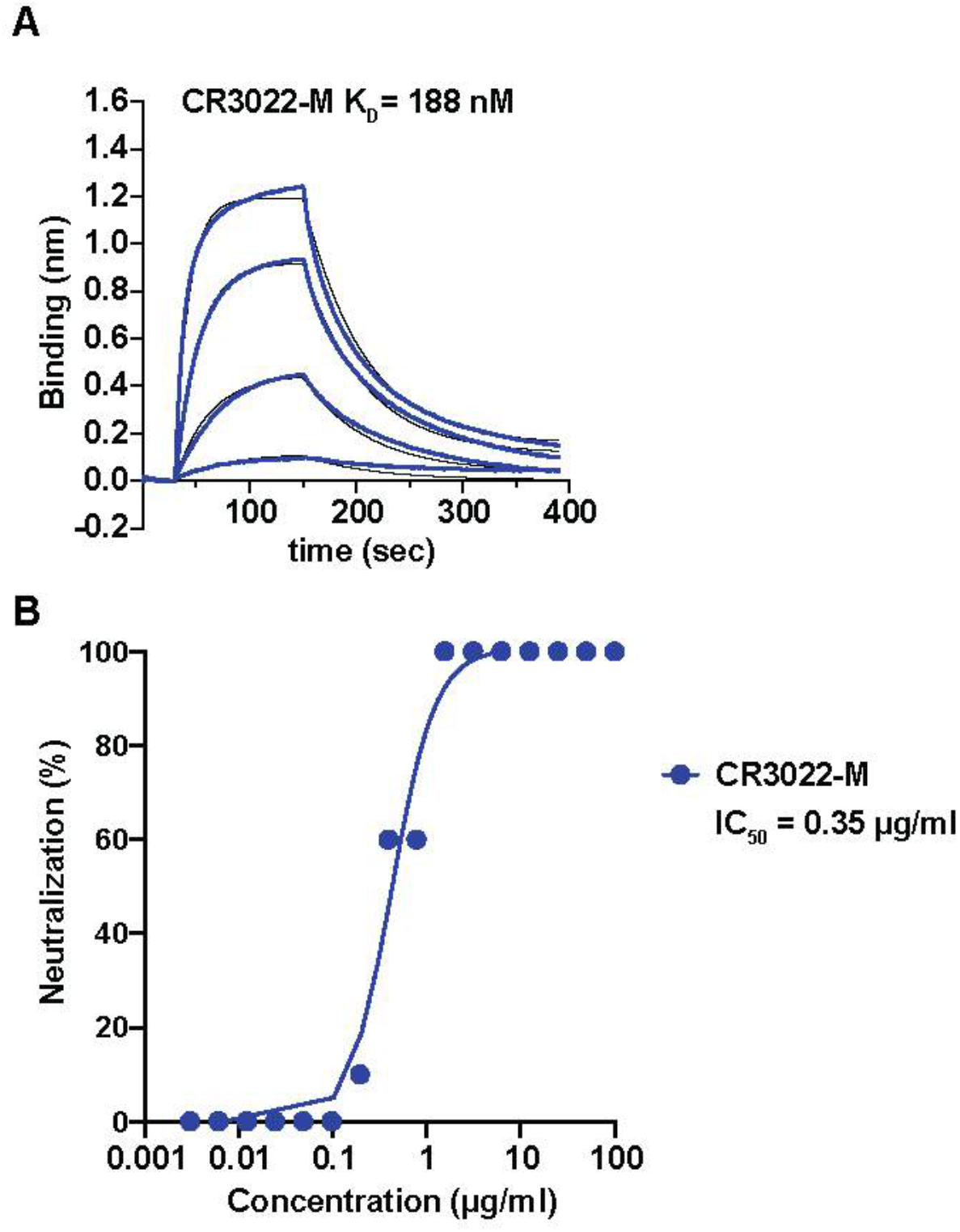
Soluble ELISA of 80R site-directed mutagenesis clones C9, D1 and D10 binding to SARS-CoV-2 (scFv antibody fragment format). Soluble scFv fragments were incubated with immobilized SARS-CoV-2 RBD on streptavidin-coated ELISA plates. Binding was detected by measuring absorbance at 450 nm.

**Supplementary Figure 5.**
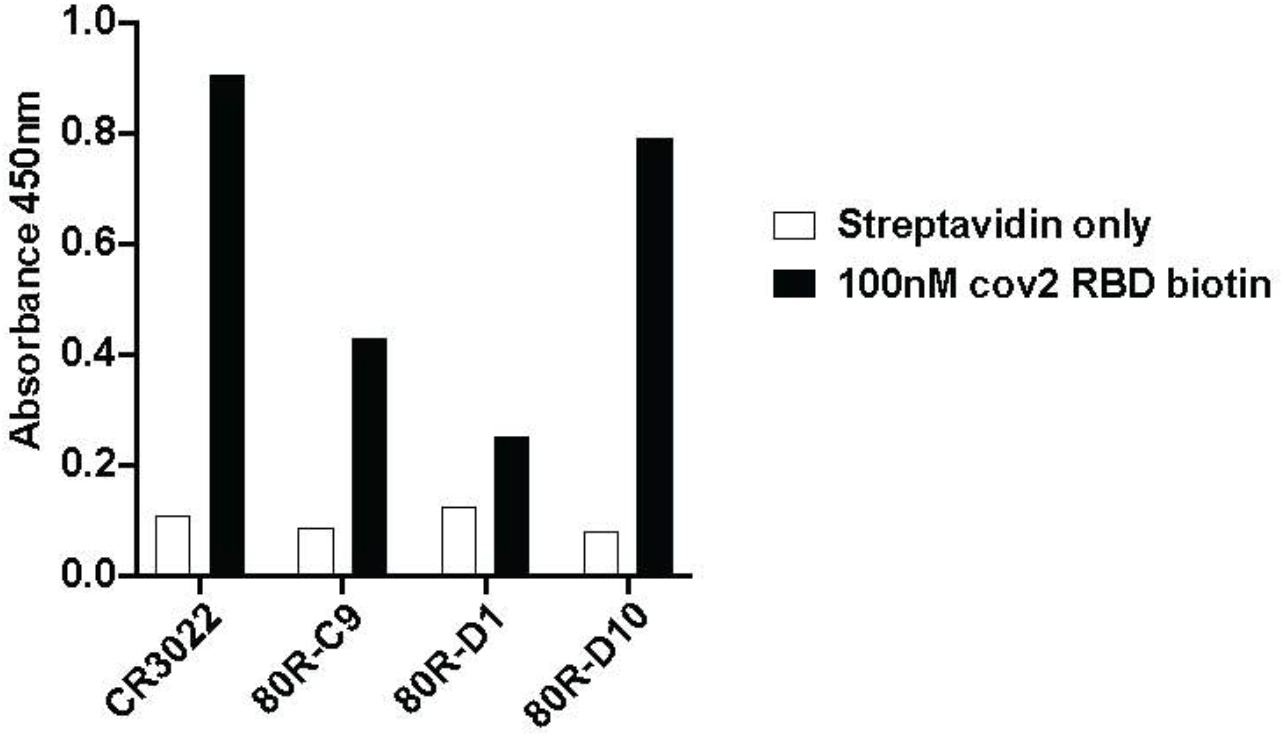
Affinity maturation of CR3022-M. **(A)** BLI affinity measurements: Biotinylated IgG was immobilized onto streptavidin sensors and incubated with different concentrations of SARS-CoV-2 RBD (from 400 nM to 50 nM by 2-fold dilutions) (see Table 1); **(B)** Neutralization of SARS-CoV-2 by CR3022-M. Serial dilutions of IgG and SARS-CoV-2 were pre-incubated for 1h at 37°C and then incubated with Human Vero E6 cells for 3 days.

**Supplementary Figure 6.**
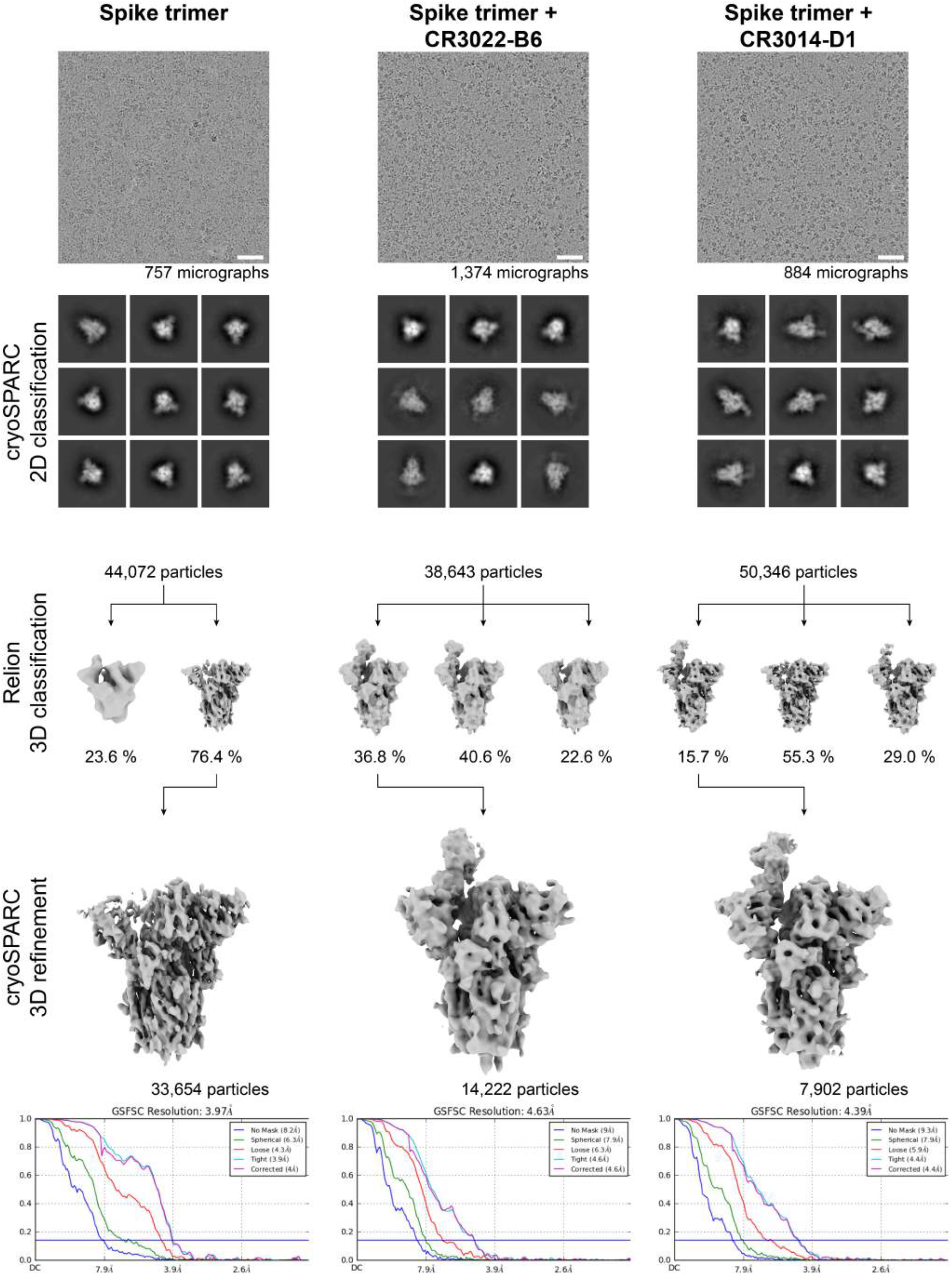
Cryo-EM data processing. Sample micrographs and flowchart to describe the data processing of the spike timer alone, or after a 1 hour incubation with CR3022-B6 or CR3014-D1. **Top:** sample micrographs, with white bar equivalent to 50 nm in length, and 9 of the best 2D class averages from each data set. **Bottom:** flowchart to describe 3D classification implemented in Relion and 3D refinement in implemented cryoSPARC, along with FSC curves with gold standard resolution estimates. Spike trimer alone had a single RBD in the up conformation. Spike trimer + CR3022-B6 classified to two classes with Fab bound to an RBD and one class without Fab bound; the class containing ~37% of the particles was chosen for full 3D refinement as the density attributed to the Fab contained higher detail. Spike trimer + CR3014-D1 classified to two classes with Fab bound to an RBD and one class without Fab bound; the class containing ~16% of the particles was chosen for full 3D refinement as the density attributed to the Fab contained higher detail. The difference between the classes containing Fab within the same dataset appeared to be caused by the RDB flexing about the RDB linker.

**Supplementary Figure 7.**
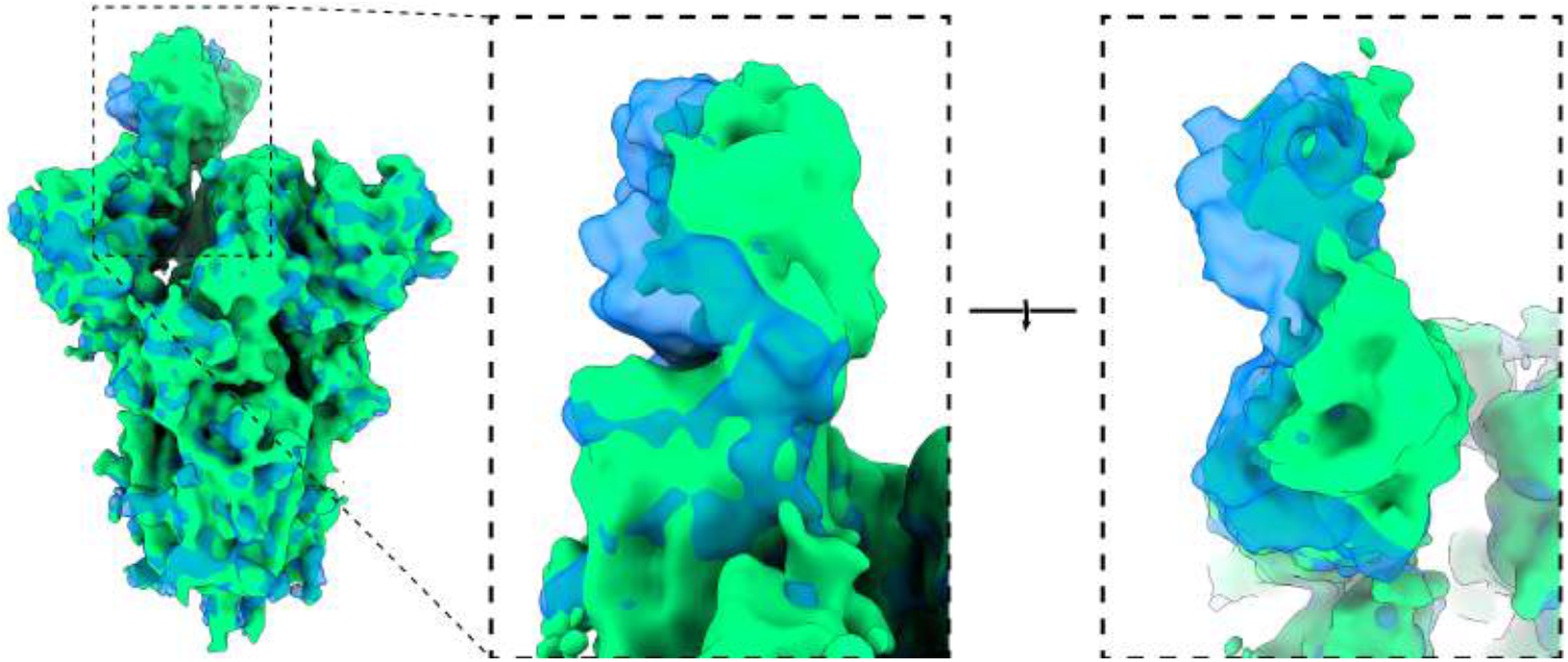
Comparison of positioning of Fab fragments in the cryo-EM structures. The trimeric spike protein in each structure was overlayed and displayed the relative orientation of the CR3022-B6 (in blue) and CR3014-D1 (in green) Fabs with each other.

**Supplementary Table 1.** List of site-directed mutagenesis diversity at CDR positions (Kabat numbering).

| Antibody CDR | Position | Codon | Encoded amino acids |
| --- | --- | --- | --- |
| CR3022-H1 | I30 | DWY | Y, D, I, N, F, V |
|  | T31 | DMY | Y, A, D, S, N, T |
| CR3022-H2 | D54 | KMT | Y, A, D, S |
|  | E56 | RMN | A, T, D, E, N, K |
| CR3022-H3 | S96 | DMY | Y, A, D, S, N, T |
|  | I98 | VYT | A, I, L, P, T, V |
|  | S99 | KMT | Y, A, D, S |
|  | T100 | DMY | Y, A, D, S, N, T |
| CR3022-L1 | Y27D | KMT | Y, A, D, S |
|  | S27E |  |  |
|  | S27F |  |  |
|  | I28 | DWY | Y, D, I, N, F, V |
|  | N29 | NHT | Y, A, D, S, N, H, I, L, F, P, T, V |
| CR3022-L2 | T53 | DMY | Y, A, D, S, N, T |
| CR3014-H1 | S30 | KMT | Y, A, D, S |
|  | D31 | KMT | Y, A, D, S |
| CR3014-H2 | R50 | BRT | Y, D, G, H, R, C |
|  | R52 | NVT | Y, A, D, S, G, H, N, P, R, T, C |
|  | N52A | DMY | Y, A, D, S, N, T |
|  | N53 |  |  |
| CR3014-H3 | G95 | KRY | Y, D, G, C |
|  | I96 | DWY | Y, D, I, N, F, V |
|  | S97 | KMT | Y, A, D, S |
|  | P98 | YMY | H, P, S, Y |
|  | F99 | KHT | Y, A, D, S, F, V |
|  | Y100 | KMT | Y, A, D, S |
| CR3014-L1 | S30 | KMT | Y, A, D, S |
|  | S31 |  |  |
| CR3014-L2 | A50 | KMT | Y, A, D, S |
|  | S53 | DMY | Y, A, D, S, N, T |
| CR3014-L3 | S91 | NRT | Y, D, S, R, N, H, C, G |
|  | Y92 |  |  |
|  | S93 |  |  |
|  | T94 | DMY | Y, A, D, S, N, T |
| m396-H1 | S31 | DMY | Y, A, D, S, N, T |
|  | T33 |  |  |
| m396-H2 | T52 | DMY | Y, A, D, S, N, T |
|  | I53 | NHT | Y, A, D, S, N, H, I, L, F, P, T, V |
|  | L54 |  |  |
|  | I56 |  |  |
|  | N58 | DMY | Y, A, D, S, N, T |
| m396-H3 | V97 | NHT | Y, A, D, S, N, H, I, L, F, P, T, V |
|  | M98 | RNS | M, A, R, N, D, E, G, I, K, S, T, V |
| m396-L1 | N27 | DMY | Y, A, D, S, N, T |
|  | S30 | KMT | Y, A, D, S |
|  | K31 | RMN | A, T, D, E, N, K |
| m396-L3 | W91 | WKG | W, L, M, R |
|  | S93 | DMY | Y, A, D, S, N, T |
|  | S94 |  |  |
| 80R-H1 | S31 | KMT | Y, A, D, S |
|  | A33 |  |  |
| 80R-H2 | Y52A | KMT | Y, A, D, S |
|  | D53 |  |  |
|  | S55 |  |  |
|  | N56 | DMY | Y, A, D, S, N, T |
| 80R-H3 | D95 | KMT | Y, A, D, S |
|  | R96 | VRV | R, H, Q, K, D, E, G, N, S |
|  | S97 | DMY | Y, A, D, S, N, T |
|  | Y98 |  |  |
| 80R-L1 | R30 | VRV | R, H, Q, K, D, E, G, N, S |
|  | S31 | DMY | Y, A, D, S, N, T |
| 80R-L2 | D50 | KMT | Y, A, D, S |
|  | S52 | DMY | Y, A, D, S, N, T |
|  | T53 |  |  |
| 80R-L3 | R91 | NDT | Y, D, S, N, R, I, L, H, G, F, C, V |
|  | W94 | WKG | W, L, M, R |

**Supplementary Table 2.**
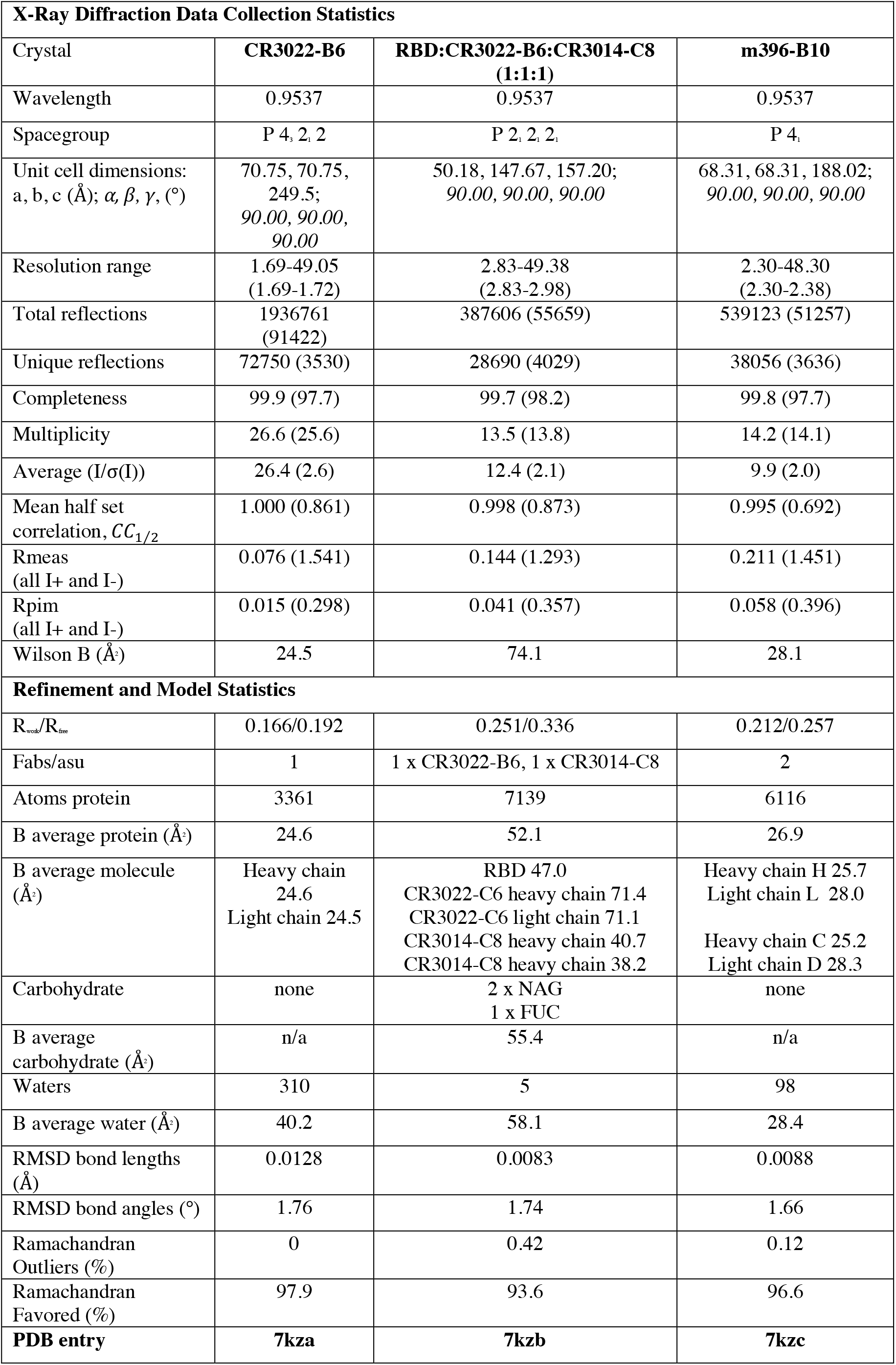
Diffraction data and model refinement statistics. Values in parentheses represent values for the highest resolution shell.

**Supplementary Table 3.** Cryo-EM data collection and refinement.

|  | #1 SARS CoV-2<br>spike trimer<br>EMD-23111 | #2 SARS CoV-2<br>spike trimer +<br>CR3022-B6<br>EMD-23112 | #3 SARS CoV-2<br>spike trimer +<br>CR3014-D1<br>EMD-23113 |
| --- | --- | --- | --- |
| **Data collection and processing** |  |  |  |
| Magnification | 150,000 | 150,000 | 150,000 |
| Voltage (kV) | 200 | 200 | 200 |
| Electron exposure<br>(e-/Å<sup>2</sup>) | 40 | 40 | 40 |
| Defocus range (μm) | 1.0-4.0 | 1.0-4.0 | 1.0-4.0 |
| Pixel size (Å) | 0.986 | 0.986 | 0.986 |
| Symmetry imposed | C1 | C1 | C1 |
| Initial particle<br>images (no.) | 44,072 | 38,643 | 50,346 |
| Final particle<br>images (no.) | 33,654 | 14,222 | 7,902 |
| Map resolution (Å)<br>0.143 FSC<br>threshold | 4.0 | 4.6 | 4.4 |
| **Refinement** |  |  |  |
| Map sharpening *B*<br>factor (Å<sup>2</sup>) | 97 | 107 | 76 |

## Supplementary Sequences

### m396-B10 (VH/VL)

- QVQLQQSGAEVKKPGSSVKVSCKASGGTFSTYSISWVRQAPGQGLEWMGGIAPSHGFANYAQKFQGRVTITTDESTSTAYMELSSLRSEDTAVYYCARDTATGGMDVWGQGTTVTVSS
- SYELTQPPSVSVAPGKTARITCGGNNIGSKSVHWYQQKPGQAPVLVVYDDSDRPSGIPERFSGSNSGNTATLTISRVEAGDEADYYCQVWDTYSDYVFGTGTKVTVL

### m396-C4 (VH/VL)

- QVQLQQSGAEVKKPGSSVKVSCKASGGTFSAYSISWVRQAPGQGLEWMGGIAPSHGTANYAQKFQGRVTITTDESTSTAYMELSSLRSEDTAVYYCARDTVTGGMDVWGQGTTVTVSS
- SYELTQPPSVSVAPGKTARITCGGNNIGYKSVHWYQQKPGQAPVLVVYDDSDRPSGIPERFSGSNSGNTATLTISRVEAGDEADYYCQVWDYTSDYVFGTGTKVTVL

### CR3022-B11 (VH/VL)

- QMQLVQSGTEVKKPGESLKISCKGSGYGFITYWIGWVRQMPGKGLEWMGIIYPGDSETRYSPSFQGQVTISADKSINTAYLQWSSLKASDTAIYYCAGGSGISTPMDVWGQGTTVTVSS
- DIQLTQSPDSLAVSLGERATINCKSSQSVLSDSIAKNYLAWYQQKPGQPPKLLIYWASSRESGVPDRFSGSGSGTDFTLTISSLQAEDVAVYYCQQYYSTPYTFGQGTKVEIK

### CR3022-G11 (VH/VL)

- QMQLVQSGTEVKKPGESLKISCKGSGYGFNYYWIGWVRQMPGKGLEWMGIIYPGDSETRYSPSFQGQVTISADKSINTAYLQWSSLKASDTAIYYCAGGDGVSTPMDVWGQGTTVTVSS
- DIQLTQSPDSLAVSLGERATINCKSSQSVLSYAVHKNYLAWYQQKPGQPPKLLIYWASTRESGVPDRFSGSGSGTDFTLTISSLQAEDVAVYYCQQYYSTPYTFGQGTKVEIK

### CR3022-B6 (VH/VL)

- QMQLVQSGTEVKKPGESLKISCKGSGYGFITYWIGWVRQMPGKGLEWMGIIYPGDSETRYSPSFQGQVTISADKSINTAYLQWSSLKASDTAIYYCAGGSGISTPMDVWGQGTTVTVSS
- DIQMTQSPSSLSASVGDRVTITCRASQSIYSALNWYQQKPGKAPKLLIYAASALQSGVPSRFSGSGSGTDFTLTISSLQPEDFATYYCQQTDIHPYTFGQGTKVEIK

### 80R-B6 (VH/VL)

- EVQLVQSGGGVVQPGKSLRLSCAASGFAFSSYAMHWVRQAPGKGLEWVAVISYDGSNKYYADSVKGRFTISRDNSKNTLYLQMNSLRAEDTAVYYCARDRSYYLDYWGQGTLVTVSS
- DIQMTQSPSSLSASVGDRVTITCRASQDIAYALNWYQQKPGKAPKLLIYAASDLQSGVPSRFSGSGSGTDFTLTISSLQPEDFATYYCQQGYKIPGTFGQGTKVEIK

### CR3014-C8 (VH/VL)

- EVQLVESGGGLVQPGGSLRLSCAASGFTFSDHYMDWVRQAPGKGLEWVGRTRNKANSYTTEYAASVKGRFTISRDDSKNSLYLQMNSLKTEDTAVYYCARGISPFYFDYWGQGTLVTVSS
- DIQMTQSPSSLSASVGDRVTITCRASQYIYDSLNWYQQKPGKAPKLLIYDSSYLQSGVPSRFSGSGSGTDFTLTISSLQPEDFATYYCQQSWDTPVTFGQGTKVEIK

### CR3014-D1 (VH/VL)

- EVQLVESGGGLVQPGGSLRLSCAASGFTFSDHYMDWVRQAPGKGLEWVGRTRNKANSYTTEYAASVKGRFTISRDDSKNSLYLQMNSLKTEDTAVYYCARGISPFYFDYWGQGTLVTVSS
- DIQMTQSPSSLSASVGDRVTITCRASQDIAYALNWYQQKPGKAPKLLIYASSSLQSGVPSRFSGSGSGTDFTLTISSLQPED FATYYCQQMGREPTTFGQGTKVEIK

